# Novel roles of Kinesin-13 and Kinesin-8 during cell growth and division in the moss *Physcomitrella patens*

**DOI:** 10.1101/819722

**Authors:** Shu Yao Leong, Tomoya Edzuka, Gohta Goshima, Moé Yamada

**Author notes:** The authors responsible for distribution of materials integral to the findings presented in this article in accordance with the policy described in the Instructions for Authors (www.plantcell.org) are: Moé Yamada and Gohta Goshima.

## Abstract

Kinesin-13 and -8 are well-known microtubule (MT) depolymerases that regulate MT length and chromosome movement in animal mitosis. While much is unknown about plant Kinesin-8, *Arabidopsis* and rice Kinesin-13 have been shown to depolymerise MTs *in vitro*. However, mitotic function of both kinesins has yet to be understood in plants. Here, we generated the complete null mutants in plants of *Kinesin-13* and *-8* in the moss *Physcomitrella patens*. Both kinesins were found to be non-essential for viability, but the *Kinesin-13* knockout (KO) line had increased mitotic duration and reduced spindle length, whereas the *Kinesin-8* KO line did not display obvious mitotic defects. Surprisingly, spindle MT poleward flux, for which Kinesin-13 is responsible for in animals, was retained in the absence of Kinesin-13. Concurrently, MT depolymerase activity of either moss kinesins could not be observed, with MT catastrophe inducing (Kinesin-13) or MT gliding (Kinesin-8) activity observed *in vitro*. Interestingly, both KO lines showed waviness in their protonema filaments, which correlated with positional instability of the MT foci in their tip cells. Taken together, the results suggest that plant Kinesin-13 and -8 have diverged in both mitotic function and molecular activity, acquiring new roles in regulating MT foci positioning for directed tip-growth.

**One sentence summary:** This study uncovered the roles of Kinesin-13 and Kinesin-8 in regulating microtubule dynamics for mitotic spindle formation and straight tip cell growth in the moss Physcomitrella patens

## Introduction

Microtubule (MT)-based motor proteins, Kinesins, form a large superfamily in animal and plant species (61 genes in *Arabidopsis thaliana*, 78 in *Physcomitrella patens*, 45 in *Homo sapiens*, and 25 in *Drosophila melanogaster*) (Reddy and Day, 2001; Miki et al., 2005; Shen et al., 2012). Kinesins show various activities in association with MTs and play pivotal roles in eukaryotic cells, such as cargo transport, MT organisation, MT dynamics regulation, and force generation (Walczak and Heald, 2008; Hirokawa et al., 2009). Comprehensive functional analysis in several animal model systems, such as fly and human cell lines, frog egg extracts, and mouse, together with biochemical characterisation of each kinesin motor have provided insights into how MT-based intracellular processes are driven and regulated during cell proliferation, differentiation, and animal development. In contrast, cellular and developmental function of plant kinesins are less clear, partly due to the difficulty in making and characterising the phenotypes of complete knockout (KO) lines of paralogous kinesins that likely function redundantly. The use of high-resolution live microscopy, which was particularly critical for assessing kinesin functions during mitosis in animals, has also been limited in plants.

Within the kinesin superfamily, Kinesin-13 and Kinesin-8 commonly show a unique activity *in vitro*: MT depolymerisation (Desai et al., 1999; Howard and Hyman, 2007; Walczak et al., 2013). *In vivo*, Kinesin-13 is able to depolymerise relatively stable MTs from both ends while MT catastrophe inducing activity is limited to the plus-end (Rogers et al., 2004; Mennella et al., 2005). The best-studied Kinesin-13, human KIF2C/MCAK, accumulates at MT ends either by diffusion or through recruitment by other MAPs (Lee et al., 2008). At the ends, it binds to and stabilises protofilament bends, which promotes strain on the association between protofilaments within the MT lattice (Moores et al., 2002; Ovechkina et al., 2002; Ogawa et al., 2017). In the mitotic spindle, KIF2C/MCAK is localised to the kinetochore and likely triggers depolymerisation of MT plus-ends that are erroneously attached to the kinetochore (Kline-Smith et al., 2004; Walczak et al., 2013). Another Kinesin-13 (human KIF2A, fly KLP10A) localises at the pole region and depolymerises MTs, including relatively stable kinetochore-bound MTs, from the minus-end, driving poleward movement of MTs (called spindle MT poleward flux) and chromosome segregation at anaphase (Rogers et al., 2004; Ganem et al., 2005; Walczak et al., 2013). Depletion of Kinesin-13 causes various mitotic errors, such as spindle elongation, spindle monopolarisation, erroneous kinetochore-MT attachment, and chromosome lagging at anaphase (Walczak et al., 2013).

While Kinesin-13 does not show motility on MTs, Kinesin-8 possesses both MT depolymerising activity and processive plus-end directed motility, and thus preferentially destabilises MT plus-ends (Howard and Hyman, 2007; Walczak et al., 2013). During mitosis, Kinesin-8 concentrates at the outer kinetochore region and prevents excessive elongation of kinetochore MTs and stabilises this kinetochore-MT attachment to promote chromosome alignment to the spindle equator (Mayr et al., 2007; Stumpff et al., 2008; Stumpff et al., 2012; Edzuka and Goshima, 2019). As a whole, Kinesin-13 and -8 MT depolymerisation activity is generally required for proper MT length regulation and correct chromosome movement during mitosis of various animal cell types (Walczak et al., 2013). This mitotic activity extends to cytokinesis, where Kinesin-13 and -8 also control anaphase spindle length and bundling, respectively (Gatt et al., 2005; Uehara et al., 2013). Kinesin-13 and -8 are also repurposed for interphase where mouse KIF2A (Kinesin-13) suppresses excessive axonal outgrowths (Homma et al., 2003), and KIF24 (Kinesin-13) and KIF19 (Kinesin-8) have roles in regulating primary cilia formation and cilia length, respectively (Kobayashi et al., 2011; Niwa et al., 2012).

Despite protein conservation, the function and activity of Kinesin-13 and -8 are not fully understood in plants. Neither mutant phenotypes nor biochemical activity have been reported for Kinesin-8. On the other hand, rice and *Arabidopsis* Kinesin-13s have been shown to preserve some degree of MT depolymerisation activity *in vitro* and *in vivo* (Oda and Fukuda, 2013; Deng et al., 2015). In *Arabidopsis thaliana* xylem vessel elements, Kinesin-13A is essential to create MT deficient areas in the cortical MT network that is utilised as a scaffold for cellulose synthase movement. Cellulose is deposited only at areas with patterned MTs, thus creating cellulose -lacking regions in MT deficient areas, called pits, allowing for lateral transport of solutes and liquids in the plant. Knockdown of Kinesin-13A by RNAi results in loss of MT patterning and smaller secondary cell wall pit formation (Oda and Fukuda, 2013). Rice Kinesin-13A was shown to be important in regulating MT dynamicity and organisation of the cortical MT network in a variety of cell types (Deng et al., 2015). However, potency of the depolymerisation activity is uncertain, since plant Kinesin-13 lacks a domain required for the robust activity of animal Kinesin-13 (Ovechkina et al., 2002; Lu et al., 2005) (Figure 1A comparing animal and plant domains) and because overexpression of Kinesin-13A in non-xylem cells did not depolymerise MTs unless coexpressed with an additional binding partner MIDD1 (Oda and Fukuda, 2013). On the other hand, Kinesin-13’s function during mitosis is unknown as the *Kinesin-13A* mutants in *Arabidopsis* and rice did not show mitotic defects. Moreover, *Arabidopsis* Kinesin-13s have been suggested to be functionally redundant as complete null mutants were embryonic lethal (Fujikura et al., 2014).

**Figure 1:**
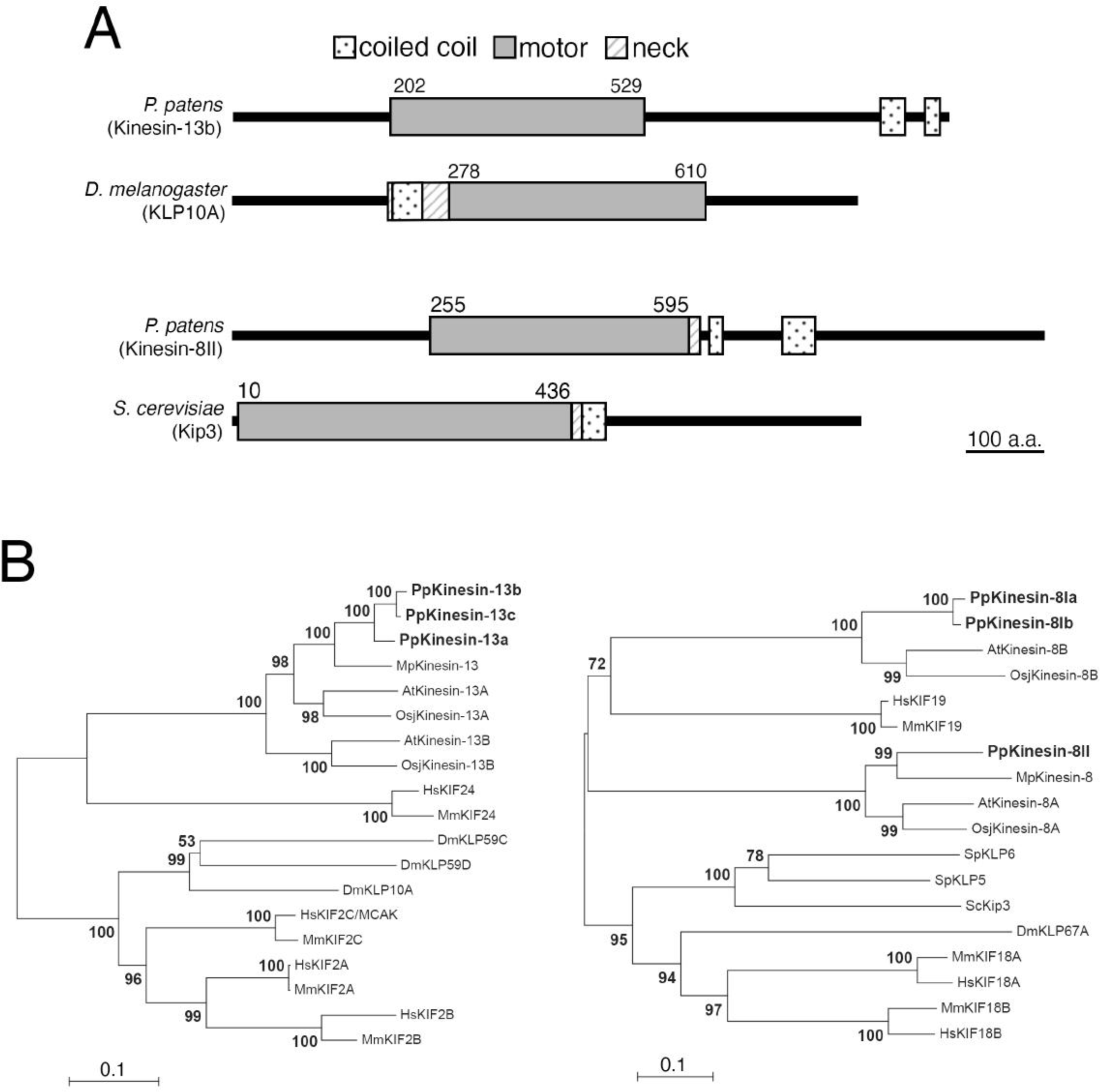
Conservation of Kinesin-13 and Kinesin-8 in the moss *Physcomitrella patens*. (A) Protein domains of Kinesin-13 (represented with Kinesin-13b) and Kinesin-8 (represented with Kinesin-8II) of moss, compared against *Drosophila melanogaster* KLP10A/Kinesin-13 and budding yeast *Saccharomyces cerevisiae* Kip3/Kinesin-8. Domains of *Drosophila* and budding yeast proteins were referenced from UniProt, whereas moss protein domains were predicted using InterPro. (B, C) Kinesin-13 and -8 phylogeny across the moss *Physcomitrella patens*, the Brassicaceae *Arabidopsis thaliana*, the liverwort *Marcantia polymorpha*, the rice *Oryza sativa* subspecies *Japonica*, the fruit fly *Drosophila melanogaster*, mammalians *Mus muculus* and *Homo sapiens*, and also for Kinesin-8 in budding yeast *Saccharomyces cerevisiae* and fission yeast *Schizosaccharomyces pombe*. After amino acid sequences were aligned with MAFFT, gapped regions were removed manually using MacClade. The phylogenetic tree was constructed using neighbour-joining methods using the Molecular Evolutionary Genetics Analysis (MEGA) software. Reliability was assessed with 1,000 bootstrapping trials. Protein sequences are presented in supplemental dataset. Horizontal branch length is proportional to the estimated evolutionary distance. Bar, 0.1 amino acid substitution per site.

In the present study, the moss *Physcomitrella patens*, a model basal plant system, was used to investigate Kinesin-13 and -8 function in general cellular processes, such as cell division. Using homologous recombination and CRISPR gene editing techniques, all three paralogues of Kinesin-13 and -8 were knocked out, generating viable complete null mutants for each of the kinesin subfamilies. We demonstrated that Kinesin-13 has a mitotic role in plants with *Kinesin-13* triple KO line having longer prometaphase duration. However, spindle MT flux was still observed and shorter metaphase spindles than the control were formed in the KO lines. In contrast, *Kinesin-8* triple KO line did not display mitotic phenotypes. Unexpectedly, neither kinesin was shown to actively depolymerise MTs *in vitro*; Kinesin-13 motor domain was able to induce MT catastrophe, while gliding activity of the Kinesin-8 motor domain was confirmed. Notably, both KO lines had wavy protonema filaments, which correlated with the MT foci abnormally fluctuating at the cell tip. Taken together, functional analyses of *Kinesin-13* and *Kinesin-8* KO in moss revealed a divergence in mitotic function and molecular activity, while revealing a novel role in regulating MT positioning for directed tip-growth.

## Results

### Kinesin-13 affects protonema growth, but not gametophore morphology

To investigate Kinesin-13’s role in the moss *Physcomitrella patens,* all three paralogous *Kinesin-13* genes (*Kinesin-13a*, *-13b*, *-13c*) (Figure 1B) were sequentially deleted by homologous recombination mediated gene replacement in the moss lines expressing GFP-tubulin and histoneH2B-mRFP (Figure S1A, B). *Kinesin-13* single and double KO moss colonies did not have observable developmental defects. Moreover, *Kinesin-13* triple KO lines (hereafter *Kinesin-13* KO) were successfully generated, indicating that *Kinesin-13s* are not essential genes in moss. There was an overall reduction in colony size in the *Kinesin-13* KO when compared to the control (Figure 2A, 2B). However, the overall morphology of the protonema colonies, gametophore (leafy shoots encasing gametangia), and rhizoids (root-like filamentous cells differentiated from gametophore basal cells) (Cove, 2005; Menand et al., 2007; Kofuji and Hasebe, 2014) were indistinguishable from the control (Figure 2A, C), which differs from the case of rice *Kinesin-13A* mutant that shows small and round grains with shortened panicles and internodes of the whole rice plant (Kitagawa et al., 2010).

**Figure 2:**
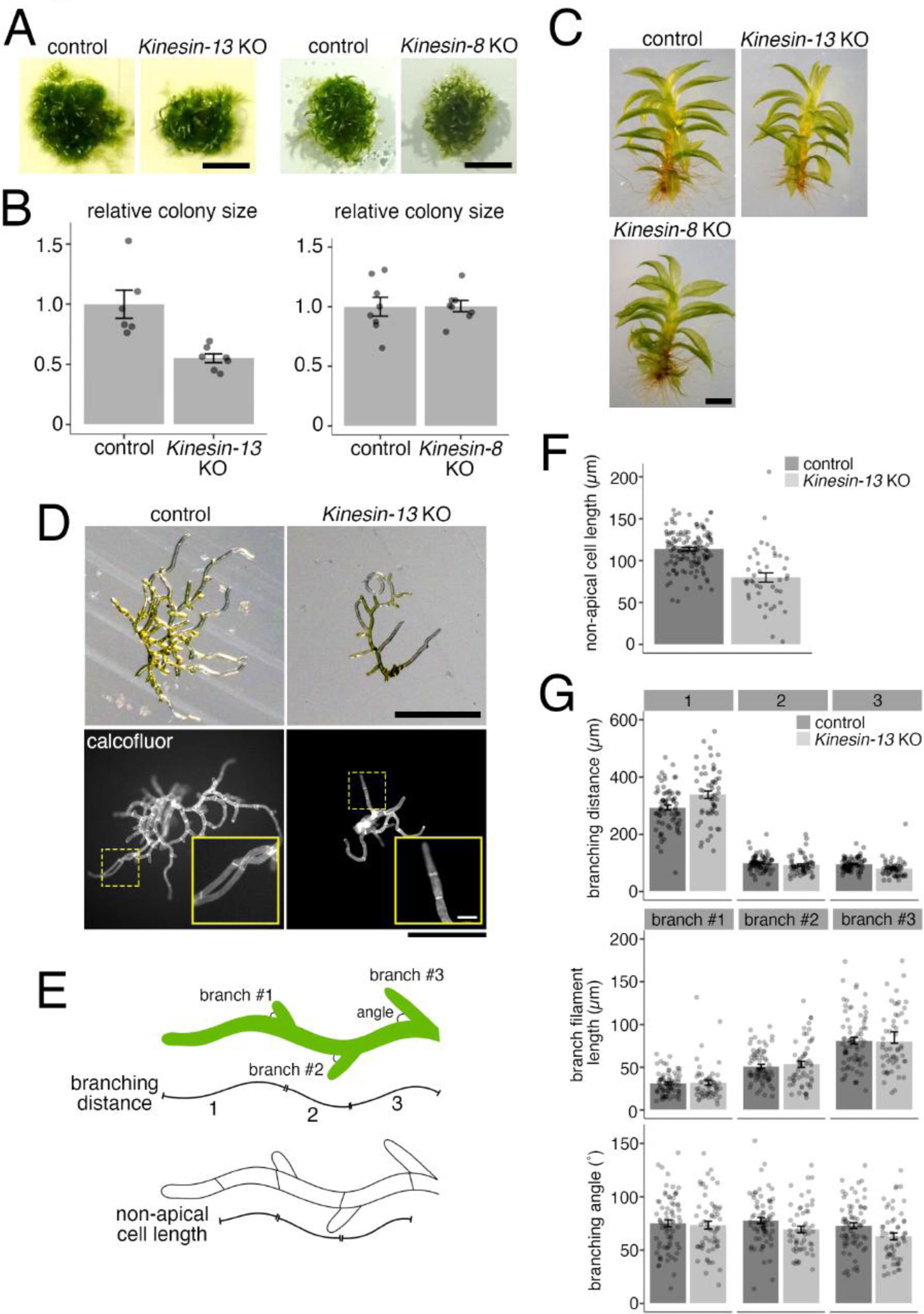
*Kinesin-13* and *-8* KO mosses are morphologically normal, but *Kinesin-13* KO moss shows retarded growth and reduced cell length. (A, B) Colony size comparison between control (*GFP-tubulin/histoneH2B-mRFP*) and *Kinesin-13* KO (GPH0438#30, left) or *Kinesin-8* KO (GPH0433#9, right) moss. Colonies were cultured from single protoplasts for 27-28 days on BCDAT where at least two independent experiments each with at least 2 plates of colonies were performed. The average colony area for each line on each plate was obtained. Actual areas were then divided by the average area of the control sample to get relative colony size. In the *Kinesin-13* KO experiment, KO moss had a relative size of 0.55 ± 0.04 (mean ± SEM; N = 7) whereas control had a relative size of 1.00 ± 0.12 (mean ± SEM; N = 6). In the *Kinesin-8* KO experiment, KO moss had a relative size of 1.01 ± 0.05 (mean ± SEM; N = 8) whereas control had a relative size of 1.00 ± 0.08 (mean ± SEM; N = 8). Points represent individual colonies, results are from one of at least two independent experiment. Bar, 5 mm. (C) Gametophore and rhizoids of control (*GFP-tubulin/histoneH2B-mRFP*) and *Kinesin-13* KO (GPH0438#6) or *Kinesin-8* KO (GPH0433#7) moss. Gametophores were isolated from colonies from small colony subcultures cultured on BCDAT for 27-28 days. Bar, 1 mm. (D) Day-8 moss colonies cultured from protoplast of control (*GFP-tubulin/histoneH2B-mRFP*) and *Kinesin-13* KO (GPH0438#30) under brightfield light (top) and with calcofluor staining (bottom). Yellow dashes boxes, inset region; bars, 500 µm; inset bar, 50 µm. (E) Cartoon depicting the measurements taken for non-apical cell length in (F) and branching phenotype analysis in (G). (F) Non-apical cell lengths of caulonema filaments were measured using calcofluor stained colonies as in (D, bottom) for control (*GFP-tubulin/histoneH2B-mRFP*) and *Kinesin-13* KO (GPH0438#30). Non-apical cell length was reduced in *Kinesin-13* KO moss to 79.9 ± 5.5 µm (mean ± SEM; N = 43), compared to control moss of 113.7 ± 1.9 µm (mean ± SEM; N = 132). Points represent individual cells; results are pooled from two independent experiments where two independent lines were analysed. (G) Branching phenotype analysis of control (*GFP-tubulin/histoneH2B-mRFP*) and *Kinesin-13* KO (GPH0438#30). In particular, branching distance of the first branch site to cell tip (top graph, leftmost bars) was increased in *Kinesin-13* KO moss to 338.4 ± 12.9 µm (mean ± SEM; N = 55), compared to control moss of 293.1 ± 8.8 µm (mean ± SEM; N = 71). Points represent individual filaments; results are pooled from two independent experiments where two independent lines were analysed.

To further investigate the colony growth phenotype in the *Kinesin-13* KO moss, early stage moss colonies regenerated from single protoplasts cultured for 8 days were analysed for non-apical cell length and protonema filament branching pattern (Figure 2D–G). Non-apical cells, which undergo little cell expansion after cell division, were found to be shorter in the *Kinesin-13* KO moss caulonema cells (Figure 2E, F), consistent with reduced cell length in rice *Kinesin-13A* mutants (Deng et al., 2015). The branching pattern was analysed by measuring the parameters of branching distance (distances from tip of protonema filament to the first three branching sites), branch filament length, and branch angle (Figure 2E). While the first branching distance (distance from tip of protonema filament to nearest branching site) increased in the *Kinesin-13* KO line, other branching pattern parameters were not observably different from that of the control (Figure 2G).

### Kinesin-13 facilitates spindle MT organisation and chromosome alignment, but does not drive spindle MT flux

The protonema tissue propagates by concerted asymmetric cell division and tip-growth in their apical stem cells (Rounds and Bezanilla, 2013). Therefore, a reduction in colony size in the *Kinesin-13* KO moss could be attributed to a defect in either or both events. To study mitosis in the *Kinesin-13* KO moss, localisation of moss Kinesin-13s to the mitotic spindle was first confirmed. As previously reported (Miki et al., 2014), moss Kinesin-13s showed spindle localisation, most enriched at the spindle equator, with the level of expression varying amongst the three paralogues; they did not show punctate signals at the spindle pole or kinetochore like animal Kinesin-13 (Supplemental Figure 2A). Next, time-lapse imaging of moss protonema cells revealed that MT-dependent nuclear movement in prophase was abnormal in the *Kinesin-13* KO line. In the control, nuclear movement is minimal or mildly apically directed as cells undergo nuclear envelope breakdown (NEBD). In contrast, in the KO line, the nucleus displayed severe retrograde movement leading up into NEBD and often continued moving basally even during spindle establishment (Figure 3A, B). This retrograde nuclear movement was also observed in the *Kinesin-13ac* double KO lines to a lesser degree, but not in the single or *Kinesin-13ab* double KO lines (Figure 3B). Additionally, overexpression of Kinesin-13b(full-length)-Cerulean under the *EF1α* promoter complemented the retrograde nuclear movement (Figure 3C, D). However, mutant Kinesin-13b constructs in which motor activity (Kinesin-13b^RIG^-Cerulean) (Dawson et al., 2007), conserved MT depolymerisation motifs (Kinesin-13b^KVD/KEC^-Cerulean) (Shipley et al., 2004), and a conserved MT binding domain (Kinesin-13b^Loop12^-Cerulean) (Soppina and Verhey, 2014) were compromised could not restore the retrograde nuclear movement (Figure 3D). Overall, these results suggest that Kinesin-13s contribute to nuclear movement redundantly in a motor-dependent manner.

**Figure 3:**
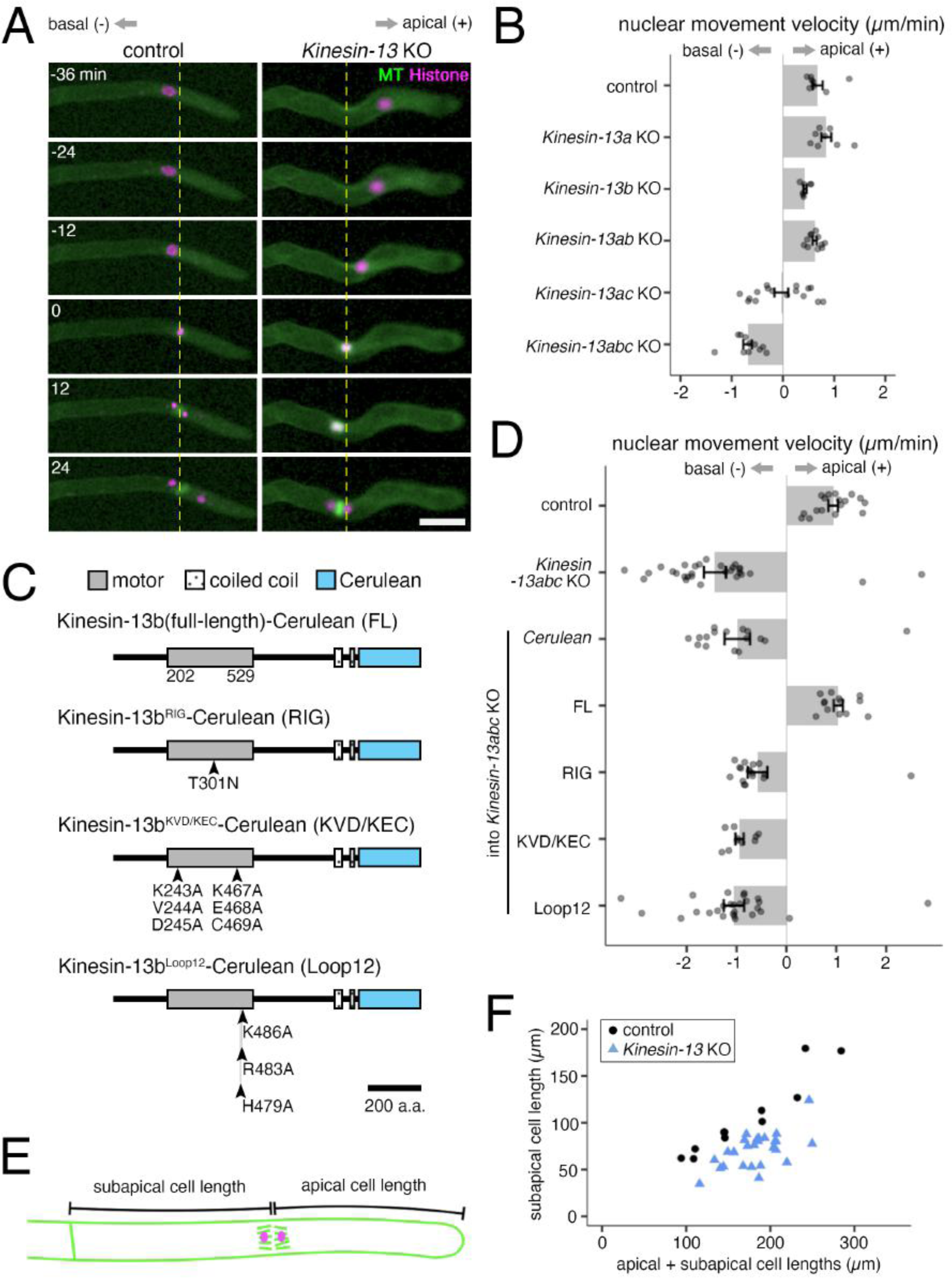
*Kinesin-13s contribute to* triple KO moss shows retrograde nuclear movement not seen in single KOs, but manifests in to a lesser degree in the *Kinesin-13ac* double KO line. (A) Snapshots of control (*GFP-tubulin/histoneH2B-mRFP*) and *Kinesin-13* KO (GPH0438#30) moss during prophase nuclear/spindle movement. *Kinesin-13* KO moss shows retrograde nuclear/spindle movement. Apical side, right, positive value for analysis in (B) and (D); basal side, left, negative values for analysis in (B) and (D); yellow dotted line, nucleus position at NEBD; bar, 50 µm; NEBD, 0 min. (B) Nuclear movement velocity during prophase of control (*GFP-tubulin/histoneH2B-mRFP*; 0.68 ± 0.10 µm/min, mean ± SEM; N = 8), *Kinesin-13a* single KO (GPH0411#43; 0.85 ± 0.10 µm/min, mean ± SEM; N = 8), *Kinesin-13b* single KO (GPH0412#11; 0.43 ± 0.03 µm/min, mean ± SEM; N = 7), *Kinesin-13ab* double KO (GPH0419#33; 0.62 ± 0.04 µm/min, mean ± SEM; N = 11), *Kinesin-13ac* double KO (GPH0420#125; -0.03 ± 0.13 µm/min, mean ± SEM; N = 16), and *Kinesin-13abc* triple KO (GPH0438#30; -0.69 ± 0.08 µm/min, mean ± SEM; N = 11). *Kinesin-13abc* triple KO shows a clear retrograde nuclear movement, whereas *Kinesin-13ac* double KO shows intermediate retrograde nuclear movement. Apically directed movement, positive values; basally directed movement, negative values. represent individual mitotic events. Results are from one of three independent experiments where two independent lines were analysed. (C) Protein domains of Kinesin-13b and mutant proteins for rescue experiments. Point mutations on Kinesin-13b-Cerulean which was introduced under the *EF1α* promoter at the *PTA1* site in the moss used for rescue experiments are illustrated. (D) Nuclear movement velocity during prophase of control (*GFP-tubulin/histoneH2B-mRFP*; 0.94 ± 0.10 µm/min, mean ± SEM; N = 17), *Kinesin-13abc* triple KO (GPH0438#30; -1.43 ± 0.22 µm/min, mean ± SEM; N = 29), *Cerulean/Kinesin-13abc* triple KO (GPH0903#1; -0.99 ± 0.25 µm/min, mean ± SEM; N = 16), *Kinesin-13b(FL)-Cerulean/Kinesin-13abc* triple KO (GPH0899#10; 1.04 ± 0.09 µm/min, mean ± SEM; N = 13), *Kinesin-13b^RIG^-Cerulean/Kinesin-13abc* triple KO (GPH0902#2; -0.58 ± 0.20 µm/min, mean ± SEM; N = 17), *Kinesin-13b^KVD/KEC^-Cerulean/Kinesin-13abc* triple KO (GPH0900#4; -0.94 ± 0.08 µm/min, mean ± SEM; N = 10), and *Kinesin-13b^Loop2^-Cerulean/Kinesin-13abc* triple KO (GPH0901#1; -1.05 ± 0.20 µm/min, mean ± SEM; N = 27). Apically directed movement, positive values; basally directed movement, negative values. Points represent individual mitotic events. Results are from one of two independent experiments where at least two independent lines were analysed. (E) Cartoon depicting how subapical and apical cell lengths were measured for (F). (F) Subapical cell length was reduced in the *Kinesin-13* KO line (GPH0438#30; 70.9 ± 3.6 µm (mean ± SEM; N = 26; p-value < 0.05)) compared to the control (*GFP-tubulin/histoneH2B-mRFP*; 105.2 ± 12.4 µm (mean ± SEM; N = 11)). Each point represents individual mitotic events. Results shown are from one of two independent experiments where two independent lines were analysed.

The severe retrograde nuclear/spindle movement during prophase likely resulted in cross cell wall positioning defects in the *Kinesin-13* KO moss. Indeed, analysis of subapical and apical cell length at anaphase onset found that subapical cell length was reduced in the *Kinesin-13* KO moss (Figure 3E, F). This correlates with reduction in non-apical cell length of early stage moss colonies, and suggests that moss Kinesin-13 has a role in cell length maintenance.

Consistent with the retrograde nuclear/spindle movement, high-resolution time-lapse imaging showed that *Kinesin-13* KO moss also has a disparity of the nucleus-surrounding MT array during prophase (Figure 4A, Movie 1). In the control, shortly before NEBD, MTs associated asymmetrically to the nucleus, with more MTs gathering on the apical side (Doonan et al., 1985; Nakaoka et al., 2012). In contrast, this apically directed MT asymmetry was altered in the KO line, with the GFP-tubulin intensity ratio of apical to basal hemispheres of the nucleus decreasing from ∼1.2 in the control to ∼1.0 (Figure 4B, C), suggesting that Kinesin-13s are important for MT organisation during prophase.

**Figure 4:**
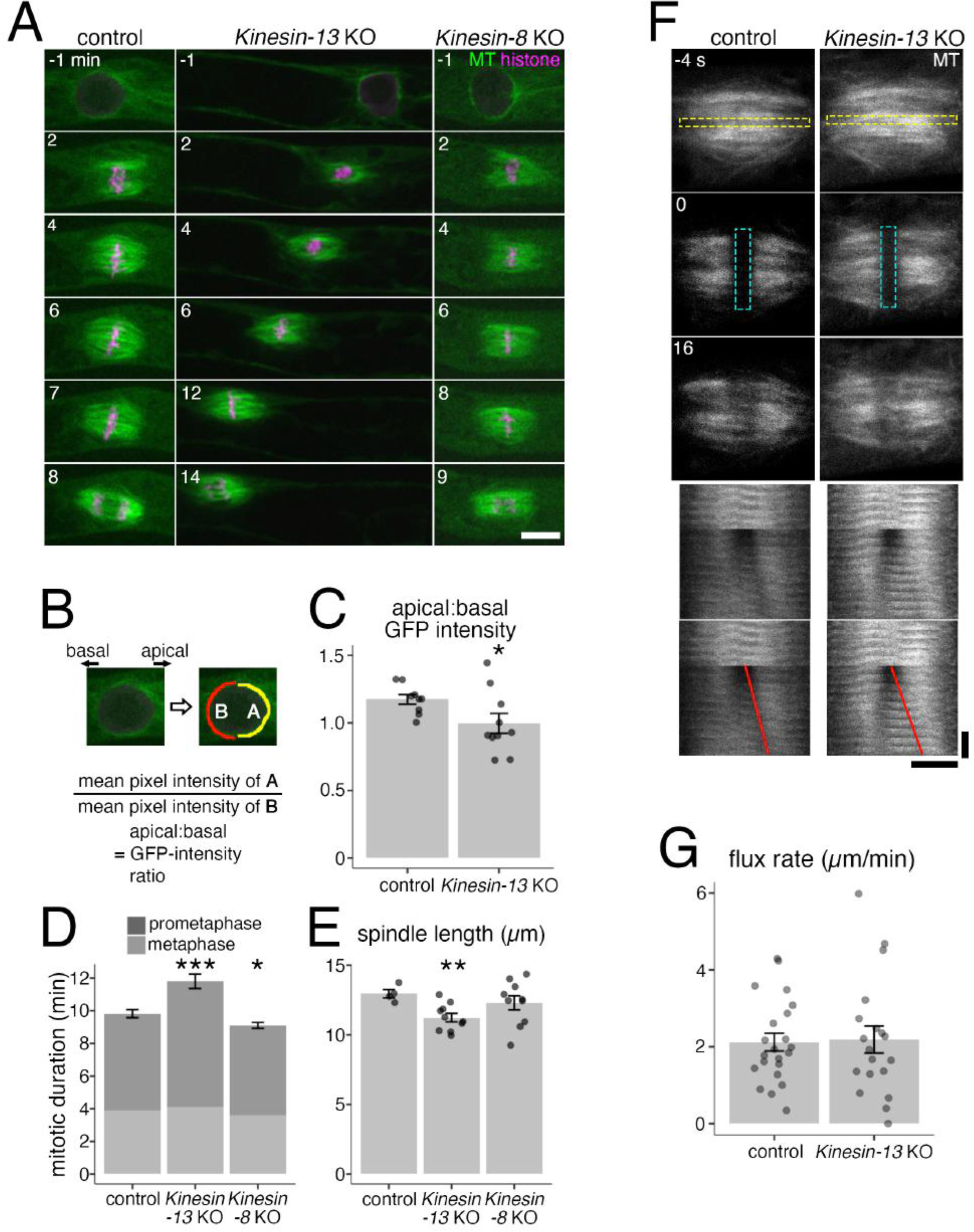
*Kinesin-13* KO moss shows defects in nuclear-proximal MT array, mitotic duration, and spindle length, but shows no difference in spindle flux rate. (A) Mitosis of control (*GFP-tubulin/histoneH2B-mRFP*), *Kinesin-13* KO (GPH0438#6), and *Kinesin-8* KO (GPH0433#9) moss. *Kinesin-13* KO showed reduced metaphase spindle length, retrograde nuclear movement during prophase, increased mitotic duration, and loss of apical bias of nuclear-MTs. *Kinesin-8* KO did not show mitotic defects. Bar, 10 μm; NEBD, 0 min; left, basal side; right, apical side. (B, C) Apical:basal GFP-intensity ratio of GFP-tubulin around the nucleus 1 min before NEBD was measured as the ratio of GFP-tubulin intensities between apical and basal hemispheric circumference. Control (*GFP-tubulin/histoneH2B-mRFP*), 1.17 ± 0.04 (mean ± SEM; N=9; p-value < 0.05); *Kinesin-13* KO (GPH0438#6, 30), 1.00 ± 0.07 (mean ± SEM; N = 10). Points represent individual mitotic events. (D) Mitotic duration of control (*GFP-tubulin/histoneH2B-mRFP*), *Kinesin-13* KO (GPH0438#6, 30), and *Kinesin-8* KO (GPH0433#7, 9) moss as measured from NEBD to anaphase onset. Control, 9.8 ± 0.3 min (mean ± SEM; N = 11); *Kinesin-13* KO, 11.8 ± 0.4 min (mean ± SEM, N = 15; p-value < 0.001). *Kinesin-8* KO, 9.1 ± 0.2 min (mean ± SEM; N = 10; p-value < 0.05). The duration of prometaphase (from NEBD to chromosome alignment) and metaphase (chromosome alignment to anaphase onset) was also measured and shown. Data shown was pooled from two independent experiments. (E) Spindle length was measured at metaphase (defined as 1 min before anaphase onset) by obtaining the distance between midpoints of apical and basal spindle widths. Control (*GFP-tubulin/histoneH2B-mRFP*), 13.0 ± 0.3 µm (mean ± SEM; N = 4); *Kinesin-13* KO (GPH0438#30), 11.2 ± 0.3 µm (mean ± SEM; N = 10; p-value <0.01); *Kinesin-8* KO (GPH0433#9), 12.3 ± 0.5 µm (mean ± SEM; N = 10). Points represent individual mitotic events. (F) Spindle poleward flux of control (*GFP-tubulin/histoneH2B-mRFP*) and *Kinesin-13* KO (GPH0438#30) moss was examined in photobleaching experiments where GFP-tubulin signals on a strip along the metaphase plate was bleached. The bleached regions separating towards the poles are indicative of spindle poleward flux function. Horizontal bar, 5 μm; vertical bar; 12 s; yellow dashed rectangle in the top panel indicates region used to make time series (bottom panel); cyan dashed rectangle represents bleached region; red lines indicate the segmented lines drawn on the kymograph to obtain flux rate in (G). (G) Quantification of spindle poleward flux experiment as shown in (F). Control, 2.1 ± 0.2 µm/min (mean ± SEM; N = 22); *Kinesin-13* KO, 2.2 ± 0.4 µm/min (mean ± SEM; N = 19). Points represent individual mitotic events, shown are results from four independent experiments.

Upon NEBD, MTs assemble into a bipolar spindle. However, spindle assembly required more time than control cells as anaphase onset was delayed, with the majority of the delay due to slow spindle MT organisation as prometaphase was delayed but metaphase was unaffected (Figure 4D). Despite the drastic nuclear movements and mitotic delay, MTs reorganised into the phragmoplast, which is the MT-based machinery required for cell plate formation, and cytokinesis was completed in 15 out of 15 cells, indicating that Kinesin-13s are dispensable in the later stages of cell division.

Unexpected from previous studies in animals and the predicted MT depolymerisation activity of Kinesin-13, the metaphase spindle was shorter, rather than longer in the KO cells (Figure 4E). In animal cells, MT depolymerisation at the spindle pole by Kinesin-13 is important for poleward flux of spindle MTs, where tubulin is flowed from the spindle equator to the pole regions through the continuous addition and removal of tubulin heterodimers at the plus- and minus-ends, respectively (Rogers et al., 2005). To investigate if Kinesin-13 depletion affects poleward MT flux in moss, GFP-tubulin at the equator of the mitotic spindle was bleached, and the movement of the photobleached strip was monitored. Surprisingly, the strip migrated towards the poles as in control cells, indicating that MT poleward flux took place in spite of complete Kinesin-13s depletion (Figure 4F, G, Movie 2). Thus, Kinesin-13 contributes to mitosis in an unconventional manner in moss.

### Kinesin-13 regulates straight growth of the protonema filament by controlling the position of MT focal points

To study the colony growth defect of the *Kinesin-13* KO line in detail, long-term time-lapse imaging of protonema filament growth was performed. Protonema filaments were found to be wavy with the protonema cell tip periodically changing growth direction in the *Kinesin-13* KO line (Figure 5A, Movie 3). The *Kinesin-13* KO line was shown to be wavier with a bend frequency of 0.024 ± 0.002 µm^-1^ (mean ± SEM; N = 26) compared to the control (0.006 ± 0.001 µm^-1^, mean ± SEM; N =28) (Figure 5B). Interestingly, the *Kinesin-13ac* double KO line showed a milder wavy phenotype, while the single and *Kinesin-13*ab double KO lines did not (Supplemental Figure 3A). Additionally, ectopic expression of full-length Kinesin-13b rescued the waviness phenotype (Figure 5B, Supplemental Figure 3B). Thus, Kinesin-13s are required for straight tip-growth.

**Figure 5:**
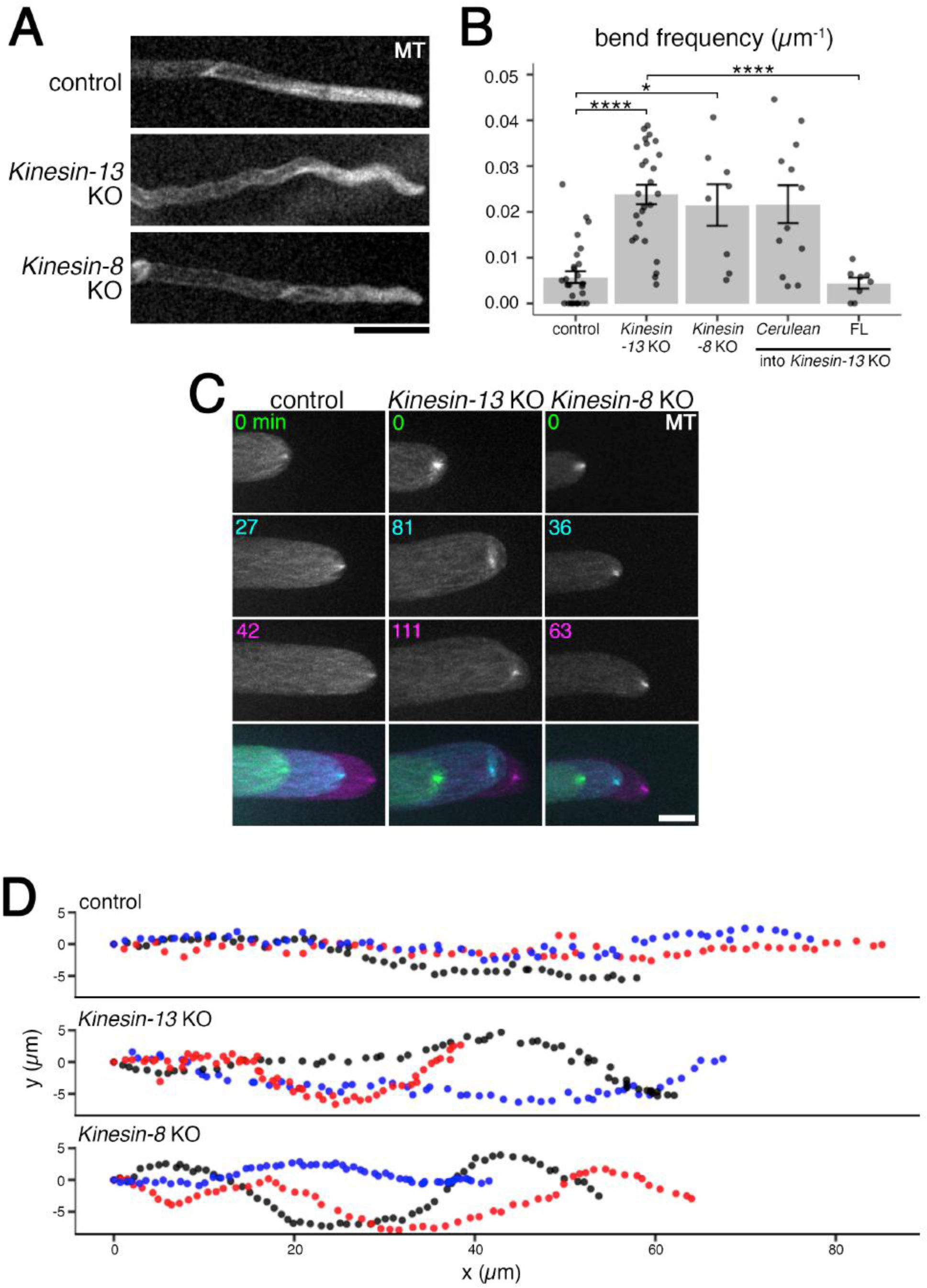
*Kinesin-13* and *-8* KO moss have wavy protonema filaments correlated with unstable MT foci positioning. (A) Protonema filaments of control (*GFP-tubulin/histoneH2B-mRFP*), *Kinesin-13* KO (GPH0438#30), *Kinesin-8* KO (GPH0433#9) moss. Bar, 50 µm. (B) Waviness of protonema filaments measured as frequency of wavy bend (>18°) of protonema filaments over measured lengths. Control (*GFP-tubulin/histoneH2B-mRFP*), 0.006 ± 0.001 μm^-1^ (mean ± SEM; N = 28 filaments); *Kinesin-13* KO (GPH0438#30), 0.024 ± 0.002 µm^-1^ (mean ± SEM; N = 26 filaments; p-value < 0.0001); *Kinesin-8* KO (GPH0433#7), 0.022 ± 0.005 µm^-1^ (mean ± SEM; N = 8 filaments; p-value < 0.01); *Cerulean/Kinesin-13* KO (GPH0903#1), 0.022 ± 0.004 µm^-1^(mean ± SEM; N = 12); *Kinesin-13b(full-length)-Cerulean/Kinesin-13* KO (GPH0899#10), 0.004 ± 0.001 µm^-1^ (mean ± SEM; N = 8; p-value < 0.0001). Points represent individual protonema filaments, results shown are from one experiment of at least four independent experiments. (C) MT foci at tip of caulonema cell of control (*GFP-tubulin/histoneH2B-mRFP*), *Kinesin-13* KO (GPH0438#30), *Kinesin-8* KO (GPH0433#9) moss. Images were acquired with z-sections at 0.3 µm intervals for 20 µm range, and maximum z-projections are displayed. Bottom panels show overlaid time series. Bar, 10 μm; colours in time series indicate different time points as labelled in top panels. (D) MT foci positions were tracked using FIJI MOSAIC plug-in 2D/3D particle tracker (Sbalzarini and Koumoutsakos, 2005) in time-lapse imaging data as in (C). (x, y) trajectories of three representative MT foci (shown in different colours) for each line are displayed. Each point represents subsequent positions at each time point, at 3 min intervals for 3 h. Same lines as in (B) are represented.

Directionality of protonema tip-growth in moss has been stipulated to be dependent on MTs (Doonan et al., 1988). At the apex of the protonema tip cell, plus-ends of MTs converge into a focus known as the MT foci (Hiwatashi et al., 2014). This occupies about the same place as the focal point of the actin filament cloud in a mutually dependent manner (Wu and Bezanilla, 2018; Yamada and Goshima, 2018). Tip-growth defects including abnormal tip branching, retarded growth, and isotropic growth are the phenotypes observed amongst transgenic mutants for regulators of cytoskeletal dynamics where its organisation at the tip is impaired (actin related proteins, myo8, KINID kinesin, KCH kinesin (Rounds and Bezanilla, 2013; Hiwatashi et al., 2014; Wu and Bezanilla, 2018; Yamada and Goshima, 2018)). As such, it is possible that *Kinesin-13* depletion may result in defective MT organisation at the cell tip, causing abnormal wavy protonema growth. MT foci behaviour in the *Kinesin-13* KO line was investigated with spinning disc confocal microscopy where the MT foci of the *Kinesin-13* KO moss was unstable and fluctuated frequently (Figure 5C, D, Movie 4). Interestingly, in 19 of 20 bending events observed, the displacement of the MT foci occurred prior to cell bending, indicating that the MT foci dictated protonema growth direction (Supplemental Figure 4). These results suggest that Kinesin-13s regulate anisotropic growth of protonema filaments by positional maintenance of the MT foci at the cell tip.

### Kinesin-13 is an interphase MT plus-end tracking protein

To investigate Kinesin-13’s localisation during interphase, endogenously tagged Kinesin-13-Citrine lines (Miki et al., 2014) was observed with spinning disc confocal microscopy. Consistent with the depletion data, Kinesin-13s localised to the MT foci (Figure 6A and Supplemental Figure 2B). To address if Kinesin-13 also associates with individual MTs in the endoplasm, we utilised oblique illumination fluorescence microscopy that enables observation of single MTs near the cell cortex with reduced effect of chloroplast autofluorescence (Jonsson et al., 2015; Nakaoka et al., 2015). In the interphase MT array, Kinesin-13s accumulated at the ends of growing MTs and disappeared from ends when MTs switched to the shrink phase (Figure 6B, C, Supplemental Figure 2C, Movie 5). Since MT minus-ends are stabilised and exhibit little to no dynamicity in this cell type (Leong et al., 2018), we concluded that Kinesin-13 localises to the plus-ends of growing MTs. The plus-end tracking behaviour is reminiscent of human KIF2C/MCAK and *Drosophila* KLP10A, which are recruited by EB1 protein to growing plus-ends (Mennella et al., 2005; Lee et al., 2008).

**Figure 6:**
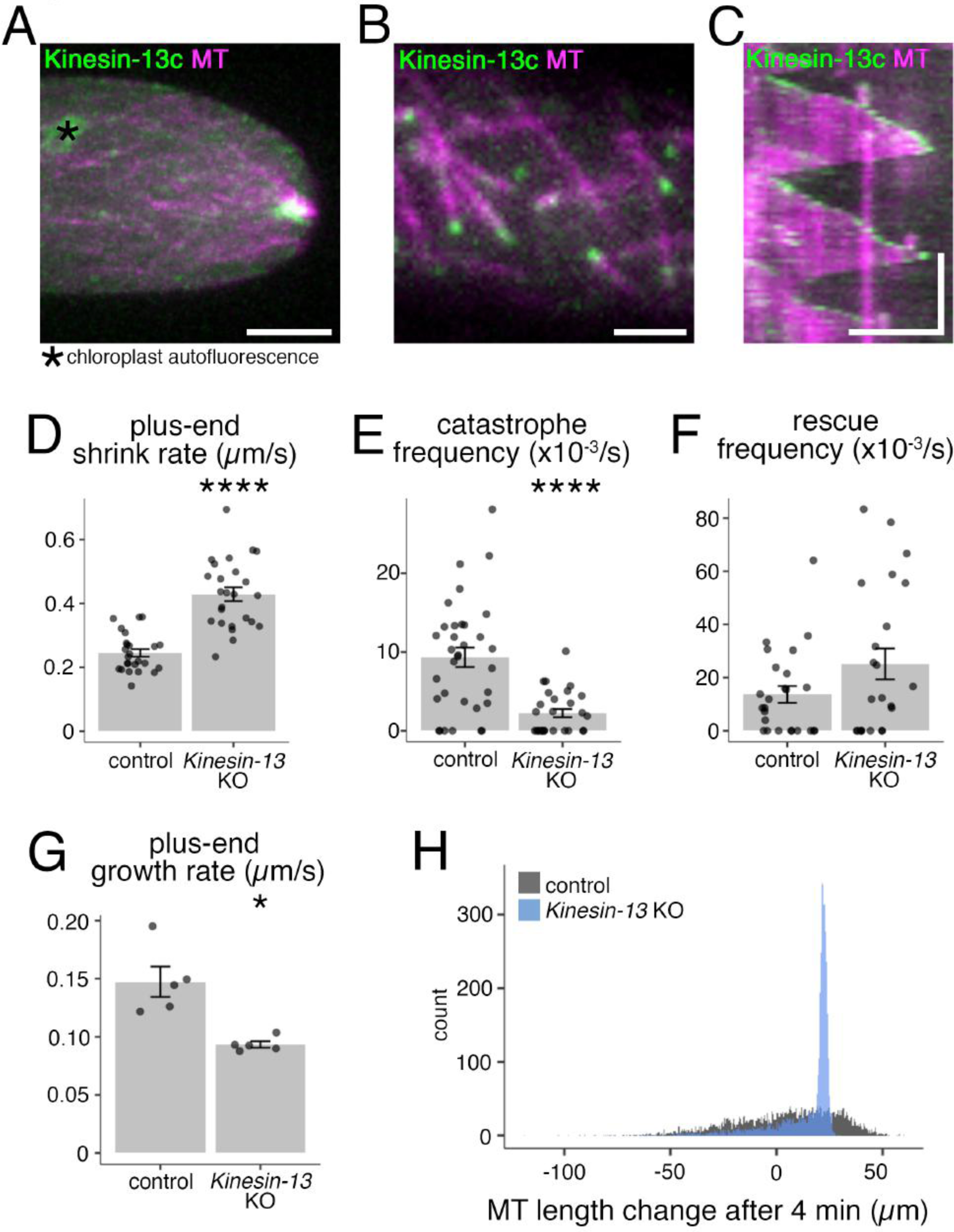
*Kinesin-13* localises to the interphase MT network and depletion of *Kinesin-13* results in increased shrink rate, reduced catastrophe frequency, increased rescue frequency, and reduced growth rate. (A) MT foci of *Kinesin-13c-Citrine/mCherry-tubulin* (GPH0100#15) moss. Image was acquired at 0.3 µm intervals for 10 µm range; shown is maximum z-projection. Bar, 5 µm. (B) Interphase MT network of *Kinesin-13c-Citrine/mCherry-tubulin* (GPH0100#15) moss. Images were acquired by oblique illumination fluorescence split-view microscopy to avoid chloroplast autofluorescence. Bar, 2 µm. (C) Kymograph of MT growth taken from imaging as in (B), taken every 3 s. Vertical bar, 2 min; horizontal bar, 5 µm. (D) Interphase MT plus-end shrink rate of control (*GFP-tubulin/histoneH2B-mRFP*, 0.245 ± 0.012 µm/s (mean ± SEM; N = 25 cells)) and *Kinesin-13* KO (GPH0438#30, 0.429 ± 0.021 µm/s (mean ± SEM; N = 25 cells; p-value < 0.0001)) moss. Points represent individual cells; results shown are from one experiment of two independent experiments. (E) Interphase MT catastrophe frequency of control (*GFP-tubulin/histoneH2B-mRFP*, 9.3 ± 1.2 x10^-3^/s (mean ± SEM; N = 33 cells)) and *Kinesin-13* KO (GPH0438#30, 2.2 ± 0.5 x10^-3^/s (mean ± SEM; N = 28 cells; p-value < 0.0001)). Points represent individual cells; results shown are from two independent experiments. (F) Interphase MT rescue frequency of control (*GFP-tubulin/histoneH2B-mRFP,* 14 ± 3 x10^-3^/s (mean ± SEM; N = 25 cells)) and *Kinesin-13* KO (GPH0438#30, 25 ± 6 x10^-3^/s (mean ± SEM; N = 23 cells)). Points represent individual cells; results shown are from two independent experiments. (G) Interphase MT plus-end growth rate of control (*EB1-Citrine/mCherry-tubulin*, GPH0379#2, 0.147 ± 0.013 µm/s (mean ± SEM; N = 5 cells, 50 MTs)) and *Kinesin-13* KO moss (GPH0577#11, 0.093 ± 0.003 µm/s (mean ± SEM; N = 5 cells, 50 MTs; p-value < 0.05)). Points represent individual cells. (H) Simulation of MT growth of 4,000 MTs in 4 min based on a probability model established using MT dynamics parameters from *in vivo* interphase MT dynamics analyses (D–G) (refer to Materials & Methods, and Table 1). Control MT dynamics parameters yielded approximately normal distributions of MT lengths and tended to have a larger population of MTs with longer lengths, with the longest 25% of MTs ranging between 23.4 to 59.8 µm in length. For MTs under *Kinesin-13* KO conditions, the distribution of MT length was narrower, with 50% of all MTs between 11.5 to 22.6 µm lengths, whereas the longest 25% of MTs ranged from 22.6 to 29.4 µm in length. Histogram bin width = 0.5 µm.

### MT shrink rate and rescue frequency increase while MT growth rate and catastrophe frequency reduce upon *Kinesin-13* depletion

Since Kinesin-13s tracked growing MT plus-ends, the effect of *Kinesin-13* deletion on MT plus-end dynamics during interphase was analysed using time-lapse oblique illumination imaging of GFP-tubulin. MT shrink rate increased upon *Kinesin-13* depletion, from 0.25 ± 0.01 µm/s (mean ± SEM; 5 MTs per cell analysed, N = 25 cells) in the control to 0.43 ± 0.02 µm/s (mean ± SEM; 5 MTs per cell analysed, N = 25 cells) in the KO line (Figure 6D). Catastrophe frequency reduced from 9.3 ± 1.2 x10^-3^/s (mean ± SEM; N = 33) in the control to 2.2 ± 0.5 x10^-3^/s (mean ± SEM; N = 28) in the KO line (Figure 6E), while rescue frequency increased from 14 ± 3 x10^-3^/s (mean ± SEM; N = 25) in the control to 25 ± 6 x10^-3^/s (mean ± SEM; N = 23) in the KO line (Figure 6F). To analyse MT growth rate, *Kinesin-13* KO moss expressing EB1-Citrine (Supplemental Figure 1C), a tracker of growing MT plus-ends, was imaged with oblique illumination fluorescence microscopy. MT growth rate based on EB1-Citrine comet movement reduced from 0.147 ± 0.013 µm/s (mean ± SEM; 10 MTs per cell analysed, N = 5 cells) in the control to 0.093 ± 0.003 µm/s (mean ± SEM; 10 MTs per cell analysed, N = 5 cells) in the KO lines (Figure 6G). The results suggest that Kinesin-13 plays a role in regulating MT dynamics in the interphase MT network.

### Altered MT dynamics parameters may underlie MT length phenotypes in *Kinesin-13* KO

Depletion of MT depolymerases or catastrophe-promoting factors causes cytoplasmic MT lengthening and spindle expansion (Howard and Hyman, 2007; Goshima and Scholey, 2010). For example, in fission yeast cells lacking catastrophe-promoting factors, cytoplasmic MTs are more frequently polymerised beyond the limits of the cell, resulting in MT bending and curling (West et al., 2001). However, shorter metaphase spindle formation (Figure 4E) and the observation that the MT foci often is unable to reach the apex of the cell tip in *Kinesin-13* KO lines (Supplemental Figure 5, Movie 4) appear to be contradictory to this general rule. We reasoned that a decrease in MT growth rate and increase in shrink rate might be limiting for overall MT length, despite significant reduction in catastrophe frequency. To evaluate this idea, we built a probability model fixed by the parameters of MT growth rate, shrink rate, catastrophe frequency, and rescue frequency, and ran a simulation in which 4,000 MTs exhibit dynamic instability for 4 min (Figure 6H, Table 1). With control parameters, a normal distribution was obtained where the 50% of MTs ranged from -12.4 to 23.4 µm lengths, and the longest 1000 MTs ranged between 23.4 to 59.8 µm lengths. In contrast, with MT dynamics parameters of *Kinesin-13* KO cells, a narrower normal distribution was obtained, with 50% of MTs having lengths of 11.5 to 22.6 µm and the longest 1000 MTs ranged between 22.6 to 29.4 µm lengths. Thus, the formation of shorter metaphase spindles and apex-displaced MT foci is a theoretically possible outcome, and actually a more likely outcome, associated with depletion of moss *Kinesin-13* that affects both MT catastrophe frequency and growth/shrink rate.

### Kinesin-13 motor domain induces catastrophe *in vitro*

To investigate the direct effect of moss Kinesin-13 on MT dynamics, recombinant Kinesin-13b^motor^-mGFP protein was expressed and purified from bacterial expression system (Figure 7A and Supplemental Figure 5A; full-length Kinesin-13 could not be obtained in either bacteria or insect culture cell expression system). The purified Kinesin-13b^motor^-mGFP protein was subjected to ‘binding-release’ experiments to confirm ATP hydrolysis activity: it bound to MTs in the presence of non-hydrolysable ATP analogue (AMPPNP) and dissociated from MTs upon ATP addition (Supplemental Figure 5B).

**Figure 7:**
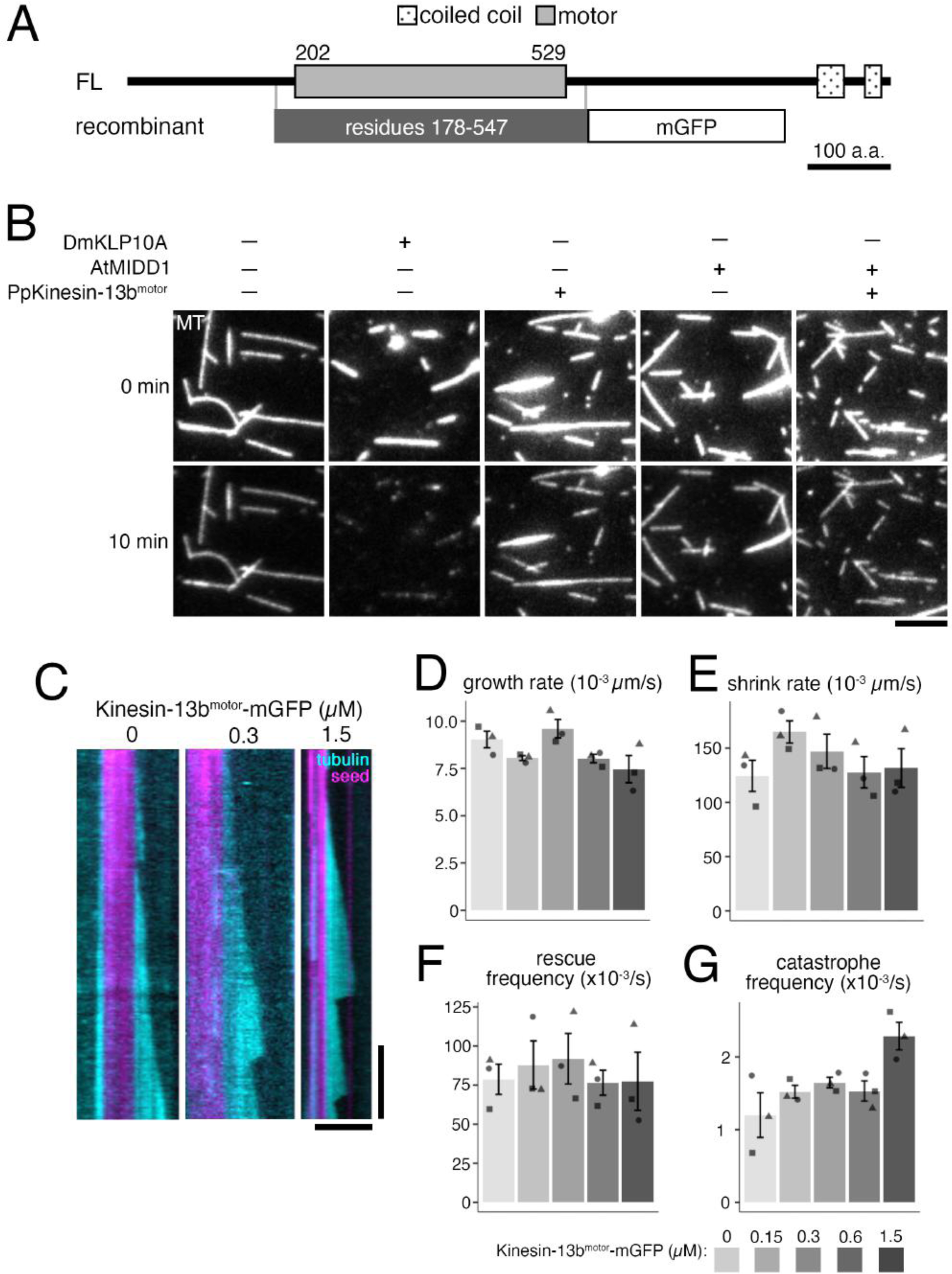
Recombinant Kinesin-13 does not depolymerise stabilised GMPCPP-MT seeds but shows MT catastrophe inducing activity. (A) Protein domains of Kinesin-13b and recombinant Kinesin-13b^motor^-mGFP construct. Protein domains were determined using InterPro. His-tag for affinity purification was attached to C-terminus of the recombinant protein. (B) *In vitro* MT depolymerisation assay using GMPCPP-stabilised MT seeds was performed using purified DmKLP10A, recombinant Kinesin-13b^motor^-mGFP construct, AtMIDD1, AtMIDD1 and Kinesin-13b^motor^-mGFP construct, and also under buffer only conditions. Only DmKLP10A successfully depolymerised MT seeds. The slight reduction in intensity in the bottom panels is due to photobleaching during imaging. All proteins were used at 200 nM except for AtMIDD1 which was at 100 nM. Bar, 5 µm. (C) Representative kymographs of *in vitro* MT dynamics polymerisation assays with Kinesin-13b^motor^-mGFP construct at 0, 0.3, and 1.5 µM. Time-lapse imaging was performed with TIRF microscopy taken every 3 s. Brightness and contrast was manually adjusted. Vertical bar, 2 min; horizontal bar; 5 µm. (D-G) *In vitro* MT dynamics parameters were analysed from time-lapse imaging of *in vitro* MT dynamics polymerisation assays with Kinesin-13b^motor^-mGFP construct at 0, 0.15, 0.3, 0.6, and 1.5 µM taken using TIRF microscopy at every 3 s. In particular, growth rate was observed to reduce slightly, from 9.0 ± 0.4 x10^-3^µm/s (mean ± SEM; N = 3) in buffer only conditions, to 7.5 ± 0.7 x10^-3^µm/s (mean ± SEM; N = 3) in 1.5 µM protein. Catastrophe frequency was observed to reproducibly increase with high concentrations of Kinesin-13b^motor^-mGFP, having a catastrophe frequency of 2.3 ± 0.2 x10^-3^/s (mean ± SEM; N = 3) at 1.5 µM protein, compared to 1.2 ± 0.3 x10^-3^/s in buffer only conditions. Points represent mean values from independent experiments.

The ATPase-active protein was added to GMPCPP-stabilised MTs, but did not show active MT depolymerisation like that of animal Kinesin-13 protein (*Drosophila* KLP10A) (Figure 7B) (Rogers et al., 2004; Moriwaki and Goshima, 2016). We considered the possibility of moss Kinesin-13 requiring a binding partner like MIDD1 for *Arabidopsis* Kinesin-13 (Oda and Fukuda, 2013). BLAST search showed that the moss does not have MIDD1 homologues, and so moss Kinesin-13 was tested for MT depolymerisation activity in the presence of *Arabidopsis* MIDD1, but also did not depolymerise MTs (Figure 7B). Overall, the purified Kinesin-13b^motor^-mGFP construct did not exhibit MT depolymerase activity under the current experimental condition..

The purified Kinesin-13b^motor^-mGFP was also subjected to an *in vitro* MT polymerisation assay at concentrations of 0, 0.15, 0.3, 0.6, 1.5 µM. While growth rate was somewhat reduced with higher Kinesin-13b^motor^ concentration (Figure 7D), shrink rate and rescue frequency were not obviously affected by Kinesin-13b^motor^-mGFP addition (Figure 7E, F). Interestingly, in the presence of Kinesin-13b^motor^-mGFP, catastrophe frequency was reproducibly increased (Figure 7G). This result is consistent with *in vivo* data in which MT catastrophe frequency decreased in the *Kinesin-13* KO line. In contrast, the recombinant protein did not reproduce the plus-end accumulation seen *in vivo*, indicating that truncated region and/or a separate factor may be required for plus-end recruitment.

### Chromosome segregation and cell division proceed normally in the absence of Kinesin-8

Mitotic phenotype associated with Kinesin-13 deletion was fairly mild. We reasoned that another MT depolymerase instead might have a major role in MT depolymerisation in mitosis of protonema filaments, and thought to investigate Kinesin-8, which shows strong depolymerisation activity during mitosis of yeast (Hildebrandt and Hoyt, 2000; Unsworth et al., 2008). To study Kinesin-8 function in the moss, all three paralogous genes phylogenetically classed into the moss Kinesin-8 subfamily (*Kinesin-8Ia, -8Ib, -8II*) (Shen et al., 2012; Miki et al., 2014) (Figure 1B) were knocked out (Supplemental Figure 1D). Moss colonies, gametophores, and rhizoids were normal in the *Kinesin-8* KO line (Figure 2A–C). High-resolution mitosis imaging did not show any defect in prophase MT organisation, spindle formation, chromosome alignment, anaphase chromosome segregation, and cytokinesis (9.1 ± 0.2 min from NEBD to anaphase onset; N = 10) (Figure 4A, D, E). We concluded that Kinesin-8s are dispensable for mitotic cell division in moss protonema filaments.

### Kinesin-8 controls positioning of the MT foci for straight tip growth

Interestingly, the *Kinesin-8* KO line also had wavy protonema filaments with bends occurring at smaller magnitudes with a bend frequency of 0.022 ± 0.005 µm^-1^ (mean ± SEM; N = 8) (Figure 5A, B, Movie 3). Tracking of the MT foci at tip cells showed that it fluctuates more frequently in the *Kinesin-8* KO line than in the control or *Kinesin-13* KO (Figure 5C, D, Movie 4), consistent with its smaller magnitudes of bends.

### Kinesin-8II^motor^ glides MT but does not show a MT depolymerisation activity *in vitro*

To analyse intrinsic activity of moss Kinesin-8, the recombinant Kinesin-8II^motor^-GFP protein was expressed and purified from bacterial expression system (Figure 8A, Supplemental Figure 5C) and was subjected to a MT depolymerisation assay. 200 nM Kinesin-8II^motor^-GFP was added to GMPCPP-stabilised MTs but could not depolymerise them, while 200 nM *Saccharomyces cerevisiae* Kinesin-8/Kip3 could depolymerise the MTs (Figure 8B). Kinesin-8II^motor^-GFP was then tested for MT gliding activity. Protein immobilised on silanised cover glass showed ability to glide GMPCPP-stabilised MTs in an ATP-dependent manner, but also could not depolymerise those MTs (Figure 8C, D).

**Figure 8:**
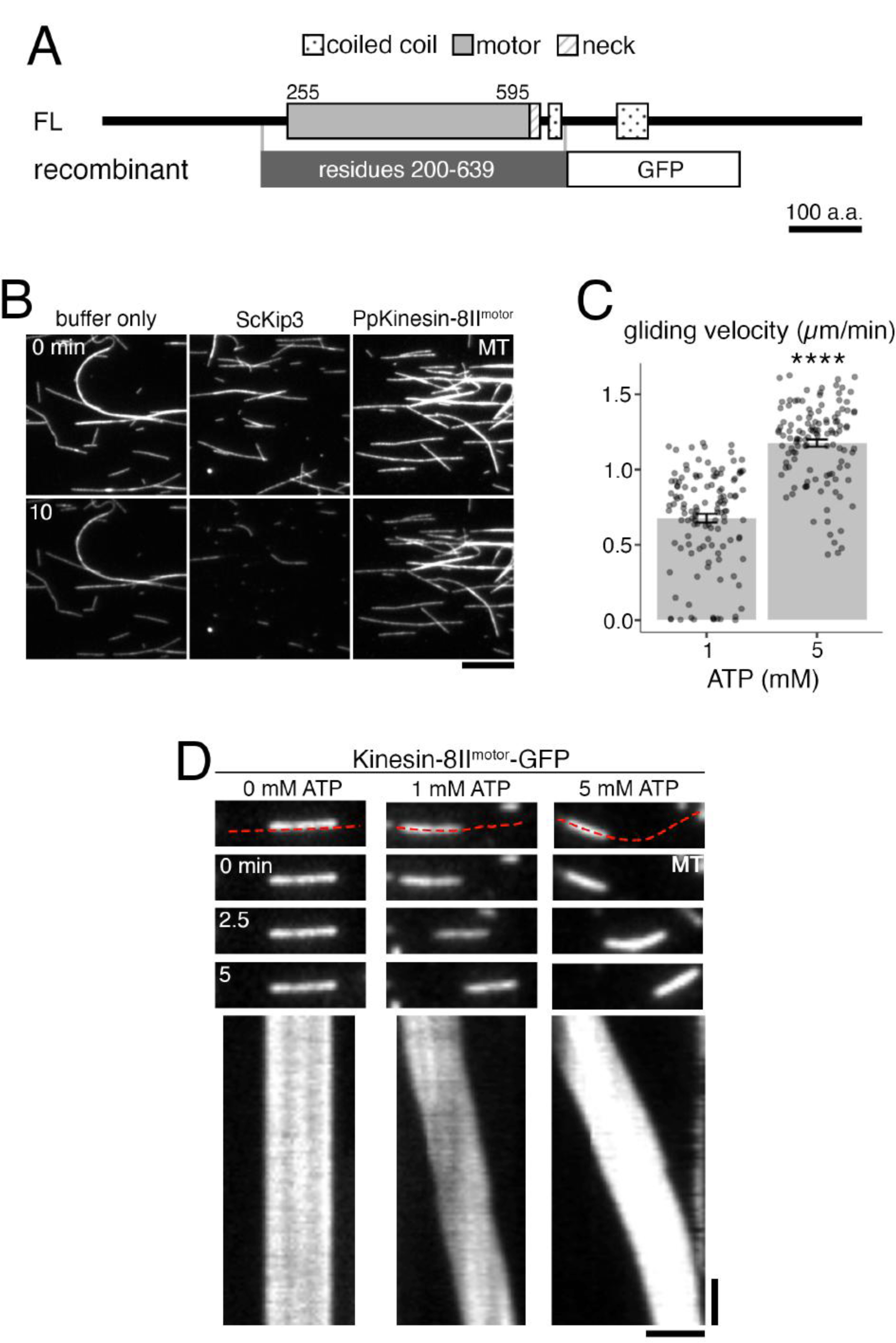
Recombinant Kinesin-8 motor does not depolymerise MTs but shows MT gliding activity. (A) Protein domains of Kinesin-8II and recombinant Kinesin-8II^motor^-GFP construct. Protein domains were identified using InterPro. His-tag for affinity purification was attached to C-terminus of the recombinant protein. (B) *In vitro* MT depolymerisation assay using GMPCPP-stabilised MT seeds was performed using purified ScKip3, recombinant Kinesin-8II^motor^-GFP, and also under buffer only conditions. Only ScKip3 showed MT depolymerisation activity. The slight reduction in intensity in bottom panels is due to photobleaching during imaging. All proteins were used at 200 nM. Bar, 10 µm. (C) ATP-dependent MT gliding velocity of Kinesin-8II^motor^-GFP. 1 mM ATP, 0.68 ± 0.03 µm/min (mean ± SEM; N = 124 MTs); 5 mM ATP, 1.18 ± 0.02 µm/min (mean ± SEM; N = 121 MTs, p-value < 0.0001). *In vitro* MT gliding assay using GMPCPP-stabilised MTs on Kinesin-8II^motor^-GFP which was immobilised on glass, at 0, 1, and 5 mM ATP. Red dotted line in top panel indicates segmented line used to draw kymographs (bottom panels). Gliding activity of Kinesin-8II^motor^-GFP was verified in 3 independent experiments. Vertical bar, 45 s; horizontal bar, 2 µm.

## Discussion

The KO lines generated in this study showed some characteristic phenotypes unreported in previous plant kinesin mutants, such as wavy cell growth accompanying MT foci positional fluctuation (Kinesin-13, Kinesin-8) and prophase MT disorganisation (Kinesin-13). Intriguingly, several processes driven by these motors in many animal and yeast species were normal in their absence in moss, such as spindle MT flux and chromosome segregation. Moreover, the hallmark activity of these kinesins, MT depolymerisation, was not detected *in vitro*. Overall, this study provides a comprehensive view on the roles of Kinesin-13 and -8 in a single plant species. Furthermore, our results reinforce the emerging view that the kinesin superfamily is well conserved in plants but have diverged in their function (Gicking et al., 2018; Nebenfuhr and Dixit, 2018).

### Are plant Kinesin-13 and -8 MT depolymerases?

Kinesin-13 is a well-known MT depolymerase in animals. *Arabidopsis* and rice Kinesin-13s have also been shown to depolymerise stabilised MTs (Oda and Fukuda, 2013; Deng et al., 2015). However, moss Kinesin-13 only exhibited catastrophe-inducing activity *in vitro* and could not depolymerise GMPCPP-stabilised MTs. This *in vitro* result is consistent with the reduced catastrophe frequency seen with interphase MTs in the *Kinesin-13* KO moss. Nevertheless, ‘negative’ results obtained *in vitro* is not necessarily conclusive: inappropriate expression systems or unsuitable biochemical environments could prevent full activity of the protein. In this study, a motor-only construct was used due to technical constraints. Thus, it is possible that other domain(s) on the *Kinesin-13* protein is required for MT depolymerisation activity. One such element may be the coiled coil, which in animal *Kinesin-13* dimerises the protein and increases MT depolymerisation activity (Hertzer et al., 2006). However, the coiled coil region required for dimerisation is located immediately upstream/downstream of the motor domain in animal Kinesin-13 (Maney et al., 2001), but is located further down the C-terminus in moss Kinesin-13 (Figure 1A); it is unclear if dimerisation of moss Kinesin-13 could enhance the activity in a similar manner to animal homologues. Furthermore, animal Kinesin-13 monomers are capable of depolymerising MTs *in vitro* (Maney et al., 2001; Hertzer et al., 2006). It is also worth noting that moss and also *Arabidopsis* Kinesin-13s lack the ‘neck’ domain that is important for strong MT depolymerisation activity in animals (Ovechkina et al., 2002); based on this feature, it was indeed originally speculated that plant Kinesin-13 might not have MT depolymerising activity (Lu et al., 2005). Thus, although it is not ruled out that moss Kinesin-13 has a MT depolymerising activity, possibly with the aid of a specific binding partner, it is enticing to say that it has diverged structurally and functionally from animal Kinesin-13.

In cells, there is even less evidence to support Kinesin-13 as a MT depolymerase. Upon *Kinesin-13* KO, interphase MTs show reduced MT growth rate and increased shrink rate. Such results instead point to Kinesin-13 being a MT growth promoter. However, MT growth promoting activity was not observed *in vitro*. This may be due to the use of the motor-only construct with which we could not recapitulate the plus-end enrichment of Kinesin-13. Alternatively, considering the decrease and increase in catastrophe and rescue frequency of interphase MTs, it is possible the Kinesin-13 regulates growth and shrink rate indirectly via tubulin cycling: reduced catastrophe would result in reduced availability of tubulin in the free tubulin pool, which might affect MT growth and shrink rates, as was proposed in the studies of *Arabidopsis* ARK proteins (Eng and Wasteneys, 2014) and more recently with plant-specific MT nucleator MACET4 (Schmidt and Smertenko, 2019).

Similar to Kinesin-13, we could not observe MT depolymerisation of the Kinesin-8 motor in our assay, which differs from human and yeast Kinesin-8. This might be due to our use of truncated construct (∼440 a.a.) as we failed to purify the longer construct (∼640 a.a.). However, we recently found that *Drosophila* Kinesin-8 (full-length) shows plus-end directed motility and induces MT catastrophe at the plus end, but is not able to depolymerise stable MTs *in vitro* (Edzuka and Goshima, 2019). Similar activities might be endowed to moss Kinesin-8.

### Kinesin-13 and -8 for mitosis

We could not detect any phenotypes in *Kinesin-8* KO lines during mitotic cell division, such as chromosome alignment and mitotic delay, which are common phenotypes observed in yeast and animal cells, suggesting that Kinesin-8 has lost mitotic functions in moss. In contrast, some but not all known mitotic functions of Kinesin-13 (Walczak et al., 2013) were observed in moss. In animal mitosis, centrosomal MTs (astral MTs) are overly developed during prophase in the absence of Kinesin-13 (Goshima and Vale, 2003; Rogers et al., 2004). Similarly, disorganised MTs were observed around the nucleus, despite the loss of centrosomes in moss (and all other land plants). Nucleus surrounding MTs may act as MTOCs equivalent to animal centrosomes. During prometaphase, kinetochore-MT attachment appears to be less efficient, since prometaphase duration was slightly prolonged; whether this was due to overall MT dynamics change or the lack of error correction, like the case of KIF2C/MCAK depletion in animal cells, remains elusive. Spindle monopolarisation that was observed in centrosome-containing animal cells (Goshima and Vale, 2003) was not detected. At metaphase, Kinesin-13 in animal cells acts as a MT depolymerase at the pole, driving MT poleward flux and halting spindle extension. Surprisingly, we could not obtain data that moss Kinesin-13 plays such a role: Kinesin-13 does not localised at the spindle pole, MT flux was detected in the *Kinesin-13* KO line, and the spindle was shorter, rather than longer, in the complete absence of Kinesin-13. MT dynamics is a major contributor to spindle length regulation in animal somatic cells (Goshima and Scholey, 2010); therefore, shortening might be due to the reduced MT growth rate observed in the endoplasm, consistent with Kinesin-13 localising at spindle equator where MT plus-ends are enriched. Chromosome segregation during anaphase A was normal, further supporting the notion that Kinesin-13 does not act as a MT depolymerase at the pole. These data indicate that the moss mitotic spindle possesses a mechanism to drive spindle MT poleward flux independent of Kinesin-13.

### Kinesin-13 and -8 for tip growth

The most prominent phenotype observed both in the *Kinesin-13* and *-8* KO lines was the tip-growth defect. Recent studies suggest the importance of the MT converging centre, the MT foci, in protonema tip-growth in moss, where F-actin, which is absolutely essential for tip-growth, is concentrated near the MT foci. In several mutants of MT-associated motors in which tip grows more slowly, the MT foci is not persistently formed (Hiwatashi et al., 2014; Wu and Bezanilla, 2018; Yamada and Goshima, 2018). These transient MT foci produced bursts of MT concentration at random locations along the tip region of the apical cell, causing the bending of the protonema filament at abrupt angles (Hiwatashi et al., 2014). Such phenotypes are similar to tip-growth defects seen when moss is treated with MT disruptive drugs (Doonan et al., 1988). Different from those mutants, we persistently observed a single MT focus in the *Kinesin-13* or *-8* KO. However, their positions were unstable, exhibiting the waviness of rather regular amplitude and frequencies. This suggests that Kinesin-13 and -8 play a role in MT foci positional guidance, rather than MT foci formation/maintenance, which ensures straight growth of the protonema filament. While straight tip-growth with limited MT foci fluctuations would allow fastest propagation of moss, wavy growth would be an advantageous mechanism to facilitate innovative exploration of the environment. Our study highlights the regulation of MT plus-end dynamics by MAPs as an intracellular mechanism to modulate cell growth in response to environmental cues, reminiscent of axon guidance in neurons (Sabry et al., 1991; Tanaka et al., 1995; Menon and Gupton, 2016).

## Materials & Methods

### Molecular cloning and gene targeting experiments

All transgenic moss lines, plasmids, and primers used in this study are listed in Table S1, S2 and S3 respectively; all lines originated from the *Physcomitrella patens* Gransden 2004 strain. Moss culture, transformation, and transgenic line selection were performed as previously described (Yamada et al., 2016). In brief, moss cells were cultured on BCDAT or BCD media under 24 h light illumination, and transformation was performed by the standard PEG-mediated method.

#### KO moss generation

*Kinesin-13* KO (GPH0438) and *Kinesin-8* KO (GPH0433) were generated in the moss strain expressing GFP-tubulin and histone-H2B-mRFP (Nakaoka et al., 2012) sequentially replacing the targeted genes with antibiotic resistance using homologous recombination (HR). To do this, 1 kb of genomic DNA sequences upstream/downstream of start/stop codons of the target genes were cloned around an antibiotic resistance cassette, and then transformed into the moss. To knock out *Kinesin-13* genes in the moss expressing EB1-Citrine and mCherry-tubulin (Kosetsu et al., 2013), *Kinesin-13b* was first removed by antibiotic resistance mediated HR as described before. For *Kinesin-13a* and *-13c*, CRISPR mediated gene removal was utilised. 20 bp guide RNAs (gRNAs) targeting the start and end of the gene were designed using CRISPOR (http://crispor.tefor.net/) (Figure S1A). gRNAs were then integrated into a BsaI site of a vector carrying U6 promoter and RNA scaffold (pCasGuide/pUC18) (Lopez-Obando et al., 2016; Collonnier et al., 2017), then the CRISPR cassettes were cloned into a vector carrying nourseothricin resistance (pSY034) with InFusion to assemble multiple gRNA cassettes into a plasmid (pSY062) following methods previously described (Leong et al., 2018). Equal amounts of this circular multicassette plasmid and *Streptococcus pyogenes* Cas9 expression vector (pGenius, (Collonnier et al., 2017)) were transformed into the *Kinesin-13b* KO/EB1-Citrine/mCherry-tubulin background. Confirmation of *Kinesin-13* and *Kinesin-8* KO lines (Figure S1B–D) were carried out by PCR as previously described (Yamada et al., 2016; Leong et al., 2018).

#### Endogenous C-terminal Citrine tagging

C-terminal endogenous Kinesin-13 and -8 Citrine tagging lines from (Miki et al., 2014) were used. In these lines, Kinesin-13 and -8 were tagged endogenously with Citrine at the C-terminal via HR where 1 kb C-terminal sequence and the downstream sequence of stop codon of the target gene flanking an antibiotic resistance cassette was used as the HR template.

Confirmation of this line was by PCR using primers designed to target the C-terminal region and outside the 3’UTR of the target gene. To make rescue plasmids, Kinesin-13 sequence was cloned from a cDNA library and ligated into the pENTR/D-TOPO vector. The Kinesin-13 mutant plasmids (Kinesin-13b^RIG^-Cerulean, Kinesin-13b^KVD/KEC^-Cerulean, Kinesin-13b^Loop12^-Cerulean) were generated by mutagenesis with the full-length Kinesin-13 plasmid and primers listed in Table S2, S3. The cloned Kinesin-13 sequences were ligated into pMN603 vector by a Gateway LR reaction, which contains EF1α promoter, *Cerulean* gene, nourseothricin-resistance cassette, *PTA1* sequences designated for homologous recombination mediated integration (Miki et al., 2016; Yoshida et al., 2019).

### Imaging sample preparation

#### Agar pad sample

Moss samples used in time-lapse imaging were prepared either in 6-well glass bottomed plates or 35 mm glass bottomed dishes as previously described (Yamada et al., 2016). Briefly, protonema cells were plated onto glasses coated with BCD agar medium and culture for 4 to 6 d.

#### Microdevice sample

Samples used for oblique illumination fluorescence microscopy were prepared in BCD liquid medium in 15 µm height PDMS microfluidic chambers mostly following previously described methods (Leong et al., 2018; Kozgunova and Goshima, 2019). Briefly, protonema cells were homogenised in BCD liquid media, filtered through a sheet of 50 µm nylon-mesh, and injected into microfluidic chambers attached unto 24 m x 32 mm micro cover glass (Matsunami, No. 1-S) and cultured for 4 to 6 days.

#### Calcofluor stained sample

8-days-old moss colonies regenerated from single protoplasts were mounted 35 mm glass bottomed dishes with BCD agar medium. The moss colonies were stained with 100 µL of mg/mL calcofluor solution diluted with water and covered with a cover glass. The samples were then imaged with Nikon Eclipse TE2000-E inverted microscope

### Moss colony assay

Protoplast regeneration assay was performed following the method described in (Vidali et al., 2007) and (Yamada et al., 2016) with some modifications. In brief, sonicated moss on cellophane-lined BCDAT plate was digested by 8% mannitol solution supplemented with 2% driselase. Generated protoplasts were washed three times with 8% mannitol solution, and 1.5 x 10^5^ mL^-1^ cells were resuspended with 7ml of protoplast regeneration liquid. After overnight incubation under the dark condition, the protoplasts were centrifuged at 510 *g* for 2 min and resuspended in 4 ml PRM solution in which CaCl_2_ was added after autoclave. 0.5–1 ml out of 4 ml protoplast solution was spread on cellophane-lined PRM plate. Then, protoplasts were cultured for 4 d and transferred to BCDAT plate, followed by 4 d culture. The 8-days-old moss colonies were observed with a stereomicroscope.

### Live *in vivo* imaging

Spinning disc confocal microscopy using 100x 1.45-NA lens and ImagEM camera (Hamamatsu Photonics) was performed with Nikon Ti inverted microscope as previously described (Yamada et al., 2016). Z-series were taken using Nano-Z Series nanopositioner combined with a Nano-Drive controller (Mad City Labs), where z-stack imaging was performed at 0.3 µm. Oblique illumination fluorescence microscopy was carried out with a Nikon Ti microscope with a TIRF unit, a 100x 1.49-NA lens, GEMINI split view (Hamamatsu Photonics), and EMCCD camera Evolve (Roper). Microscopes were controlled by Micromanager or NIS-Elements (Nikon). Low magnification epifluorescence imaging was carried out using Nikon Eclipse TE2000-E inverted microscope with 10x/0.3 LN1C Plain Fluo lens and Zyla 5.5 sCMOS camera (Andor), controlled with IQ3 (Andor). Photobleaching experiments were performed using an LSM780-DUO-NLO confocal microscopy system (Zeiss) using Plan-Apochromat 63x/1.40 Oil DIC lens controlled using Zen (Zeiss) with 489 nM diode laser (time-lapse imaging taken at 2 mW and bleaching at 100 mW). Moss expressing GFP-tubulin (GPH0438#30 for *Kinesin-13* KO and *GFP-tubulin/histoneH2B-mRFP* for control) were used for photobleaching experiments where images were acquired every 3 s. All imaging was performed at room temperature in the dark. Moss colonies and gametophores were imaged with SMZ800N (Nikon) and ILCE-QX1 camera (SONY). The stereomicroscope system was controlled with PlayMemories software (SONY).

### Computer simulations

Simulations were built in R (version 3.6.0) (https://github.com/TomoyaEdzuka/MT_dyanamics_simulation). Parameters used in this simulation for each condition (control or *Kinesin-13* KO) are listed in Table 1. In this simulation, catastrophe and rescue frequency parameters were used to determine the probability of individual steps (1 s) undergoing a transition change or to continue a growth/shrink phase. At each phase transition (i.e. catastrophe/rescue event) new growth/shrink rates were assigned following a log normal distribution of the growth and shrink parameters. MT lengths were then simulated for 4,000 trials (i.e. 4,000 MTs) for 240 steps (i.e. 4 min).

### Protein purification

The motor domain of Kinesin-13b, which is the most highly expressed protein of the three paralogous *Kinesin-13* genes, was cloned from moss cDNA and transgenically linked to monomeric GFP (mGFP) and 6xHis at the C-terminus (plasmid pGG885, Table S2). Kinesin-13b^motor^-mGFP expression was induced in SOLBL21 *E. coli* with 0.1 mM IPTG for 20 h at 18 °C. Harvested cells were lysed using the Advanced Digital Sonifier D450 (Branson) in lysis buffer (50 mM Tris-HCl [pH 8.0], 100 mM KCl, 2 mM MgCl_2_, 20 mM imidazole, 0.1 mM ATP) supplemented with 10 mM β-mercaptoethanol and protease inhibitors (1 mM PMSF and peptide inhibitor cocktail:5 mg/mL Aprotinin, 5 mg/mL Chymostatin, 5 mg/mL Leupeptin, 5 mg/mL Pepstatin A). After rotation with nickel-NTA coated beads for 2 h at 4 °C, the proteins were eluted using 500 µL elution buffer (50 mM Tris-HCl [pH 8.0], 100 mM KCl, 2 mM MgCl_2_, 300 mM Imidazole, 0.1 mM ATP). Elution was repeated 5 to 7 times. The eluates were then further purified through an AMPPNP-ATP ‘binding-release’ experiment. Eluates were first bound with 1 mM AMPPNP to 76.5 µM of 1 mM GMPCPP-stabilised MTs, and sedimented through an 80% glycerol cushion. Finally, the proteins were released from the pellet with 10 mM ATP. The motor domain and nearby coiled-coil domain (residues 200-639) of Kinesin-8II was cloned from moss cDNA and transgenically joined to GFP and 6xHis tag at the C-terminus (pTM266, Table S2), and introduced into a pET-23 *E. coli* expression vector. The recombinant protein was purified from SOLBL21 *E. coli* induced with 0.2 mM IPTG for 20 h at 18 °C. Harvested cells were lysed using the Advanced Digital Sonifier D450 (Branson) in lysis buffer (25 mM MOPS [pH 7.0], 250 mM KCl, 2 mM MgCl_2_, 5% sucrose, 30 mM imidazole, 0.1 mM ATP) supplemented with 5 mM β-mercaptothanol and protease inhibitors (0.5 mM PMSF and peptide inhibitor cocktails). After rotation with nickel-NTA coated beads (0.5 mL bed volume) for 2 h at 4 °C, proteins were eluted with 500 μL elution buffer (25 mM MOPS [pH 7.0], 250 mM KCl, 2 mM MgCl_2_, 400 mM imidazole, 5% sucrose, 1 mM ATP, 5 mM β-mercaptoethanol). Elution was repeated 5 to 7 times. Eluates were used immediately. For the *in vitro* MT depolymerisation assay, the buffer for the elute was exchanged to 1x Standard Assay Buffer (SAB: 25 mM MOPS [pH 7.0], 75 mM KCl, 2 mM MgCl_2_, 1 mM EGTA) supplemented with 1 mM ATP to remove imidazole using PD-25 column (GE Healthcare). *Drosophila* KLP10A (plasmid pGG885) was purified referencing (Moriwaki and Goshima, 2016), and was purified with the same protocol as Kinesin-13b^motor^-mGFP. Instead of the ‘binding-release’ experiment, elutes of KLP10A were subjected to buffer exchange to 1xMRB80 with 75 mM KCl and 0.1 mM ATP using a PD-25 column (GE Healthcare). AtMIDD1 was purified following (Oda and Fukuda, 2013), with some modifications. Briefly, GST-AtMIDD1 expression was induced in SOLBL21 *E. coli* using 0.2 mM IPTG, and cultured for 20 h at 18 °C. Harvested cells were lysed using the Advanced Digital Sonifier D450 (Branson) in lysis buffer (10 mM HEPES [pH 7.4], 1 mM EGTA, 1 mM MgCl_2_, 150 mM KCl) supplemented with 1 mM DTT and protease inhibitors (0.5 mM PMSF and peptide inhibitor cocktails). After rotation with Glutathione Sepharose 4B beads (GE Healthcare, 0.5 mL bed volume) for 2 h at 4 °C, proteins were eluted with 500 μL elution buffer (100 mM HEPES [pH 7.4], 100 mM KCl, 30 mM reduced glutathione). Elution was repeated 5 to 7 times. Buffer was then exchanged using a PD-10 column (GE Healthcare). Budding yeast Kip3 was purified following Kip3 purification (Gupta et al., 2006) and *Drosophila* KLP67A purification (Edzuka and Goshima, 2019). In brief, ScKip3-sfGFP-6xHis (pED273) was expressed in Sf21 moth cells at 27 °C for 72 h. Cells were lysed with 1% Triton X-100 in lysis buffer (50 mM MOPS [pH 7.0], 250 mM NaCl, 2 mM MgCl_2_, 1 mM EGTA, 20 mM imidazole, 0.1 mM ATP) supplemented with 2 mM β-mercaptothanol and protease inhibitors (0.5 mM PMSF and peptide inhibitor cocktails) for 30 min at 25 °C. After rotation with nickel-NTA coated beads (0.5 mL bed volume) for 2 h at 4 °C, proteins were eluted with 500 μL elution buffer (25 mM MOPS [pH 7.0], 250 mM KCl, 2 mM MgCl_2_, 400 mM imidazole, 5% sucrose, 1 mM ATP, 5 mM β-mercaptoethanol). Buffer was then exchanged to 1xSAB supplemented with 1 mM ATP to remove imidazole using PD-25 column (GE Healthcare).

### *In vitro* MT depolymerisation

The *in vitro* MT depolymerisation assay in (Moriwaki and Goshima, 2016) and (Gell et al., 2010) was followed with some modifications. DmKLP10A, Kinesin-13b^motor^-mGFP, AtMIDD1, and AtMIDD1 and Kinesin-13b^motor^-mGFP were mixed with 30% Alexa Fluor-568 labelled GMPCPP-MT seeds immobilised on silanised cover glass in assay buffer with (1x MRB80, 1 mM ATP, 50 mM glucose, 0.5 µg/µL κ-casein, 0.1% methylcellulose) supplemented with an oxygen scavenger system. Similarly, Kinesin-8II^motor^-GFP and ScKip3 were also introduced to immobilised GMPCPP-MT seeds, but in a different assay buffer (1xSAB, 0.1% methylcellulose, 50 mM glucose, 0.5 µg/µL κ-casein, 1 mM ATP, 75 mM KCl, supplemented with oxygen scavenger system). Proteins were used at 200 nM concentrations, except AtMIDD1 which was used at 100 nM following (Oda and Fukuda, 2013). TIRF imaging was taken every 3 s for 10 min at 25 °C.

### *In vitro* MT polymerisation assay

We largely followed methods previously described (Li et al., 2012; Moriwaki and Goshima, 2016; Leong et al., 2018) for the MT polymerisation assay. MT growth was initiated by flowing 20 µM porcine brain tubulin (containing 10% Alexa Fluor 647-labelled tubulin) and 0, 0.15, 0.3, 0.6, and 1.5 µM purified Kinesin-13b^motor^-mGFP in assay buffer (1x MRB80, 75 mM KCl, 1 mM ATP, 50 nM glucose, 1 mM GTP, 0.8 µg/µL κ-casein, 0.1% methylcellulose) supplemented with an oxygen scavenger system. TIRF imaging of Alexa Fluor-568 labelled GMPCPP-MT seeds and Alexa Fluor-647 labelled free tubulin was taken every 3 s for 10 min at 25 °C. Kymographs were drawn for 25 trackable dynamic MTs, and were analysed for catastrophe events and rescue events. Catastrophe frequency was then defined by the number of catastrophe events per growth time (min), whereas rescue frequency was defined by the number of rescue event per shrink time (min). Growth and shrink rates were taken from the corresponding slopes from the same kymographs. Three independent experiments, each with a different batch of purified proteins were performed.

### *In vitro* MT gliding assay

For the gliding assay with purified Kinesin-8II^motor^-GFP, methods described in (Miki et al., 2015) were referenced. Briefly, 10 µL of the freshly purified recombinant protein was introduced into the flow chamber and incubated at room temperature for 2 min in the dark, then washed with 20 µL 1x Standard Assay Buffer (SAB: 25 mM MOPS [pH 7.0], 75 mM KCl, 2 mM MgCl_2_, 1 mM EGTA). Then 10 µL reaction mix (1xSAB, 0.1% methylcellulose, 50 mM glucose, 0.5 µg/µL κ-casein, 50 mM GMPCPP-MT seeds, 75 mM KCl, supplemented with oxygen scavenger system and varying concentrations of ATP) was introduced into the flow chamber, and it was sealed with candle wax. TIRF imaging of Alexa Fluor-647 labelled GMPCPP-MT seeds in *in vitro* MT gliding assays with Kinesin-8II^motor^-GFP at varying ATP concentrations addition was taken every 3 s for 10 min at 23–25 °C. Kymographs were drawn for 30∼50 trackable dynamic MTs per sample, and the slopes of the kymographs were taken as gliding velocity.

### Data analysis

#### Moss colony growth rate

GPH0438#30 (*Kinesin-13* KO) and GPH0002#5 (control) protoplasts were made following (Yamada et al., 2016) with some modifications. In brief, moss was incubated with 2% driselase solution (in 8% mannitol), washed thrice with 8% mannitol, incubated overnight in protoplast liquid medium, and then plated in PRM/T medium on cellophane-lined PRM plates, cultured at 25 °C under continuous light. On day 2, moss-lined cellophane was transferred to BCDAT plates. Around day 7 when individual colonies were larger, they were picked and inoculated on BCDAT plates, and cultured at 25 °C under continuous light. On day 27–28, images of plates with grown colonies were taken with the in-built camera of Xperia X Performance (23 MP Type 1/2.3’ Exmor RS sensor, 24 mm equivalent lens with f/2.0 aperture). Images were analysed with FIJI, where images were first converted to 8-bit, automatically thresholded, and binarised. Colonies were automatically outlined with the wand tool and resulting area was obtained. Data in Figure 2B is pooled from three independent experiments.

#### Non-apical cell length

Low magnification calcofluor stained images of moss colonies were used to measure non-apical cell length (see Figure 2E cartoon), where only caulonema cells were measured. To distinguish between caulonema and chloronema cells, protonema filaments were first judged by sight in bright field images of the same colonies, in which chloroplast density was used as an indicator of cell state (caulonema, chloroplast sparse; chloronema, chloroplast dense).

#### Branching distance, branch filament length, and branching angle

Low magnification images of 8-days-old moss colonies regenerated from single protoplasts were used to analyse branching pattern parameters. Branching parameters were manually measured as shown in cartoon Figure 2E.

#### Nuclear movement velocity

Samples from epifluorescence imaging of moss protonema filaments undergoing mitosis were analysed. Kymographs were drawn on the filaments, and nuclear movement velocity was obtained from the slopes of nuclear movement in these kymographs where positive and negative values were assigned to apical and basal directions, respectively.

#### Subapical cell length

Samples from epifluorescence imaging of moss protonema filaments undergoing mitosis were analysed. Kymographs were drawn on the filaments, and the lengths from the middle of the spindle at anaphase to the cell tip and to the previous cell plate was measured as apical and subapical cell length, respectively (see Figure 3E cartoon).

#### GFP-tubulin intensity around the nucleus

Spinning disc confocal fluorescence time-lapse imaging of GFP-tubulin and histoneH2B-mRFP moss taken every 1 min was used for analysis. Segmented line built-in tool in FIJI was used to mark the hemispheric circumference around the nucleus on the apical and basal side at 1 min before NEBD (see Figure 4B cartoon) and mean pixel intensity was measured. Apical:basal GFP-intensity ratio was defined as the ratio of the mean pixel intensity of the apical hemisphere over that of the basal hemisphere.

#### Mitotic duration

Mitosis images taken every 1 min with spinning disc confocal microscope were used for analysis. Mitotic duration was defined as time from NEBD to anaphase onset, whereas prometaphase is the time from NEBD to chromosome alignment, and metaphase duration is the time from chromosome alignment to anaphase onset.

#### Spindle length

Mitosis images taken every 1 min with spinning disc confocal microscope were used for analysis. The metaphase spindle (defined as 1 min before anaphase onset) was used to analyse spindle area. 4 points (the two limits of basal and apical pole widths) were marked and their (x, y) positions were obtained, where spindle length was obtained from the distance between the midpoints of the basal and apical pole widths.

#### Spindle MT flux rate

Mitosis images were taken every 3 s with LSM780-DUO-NLO confocal microscope. *Kinesin-13* KO (GPH0438#30) and control (*GFP-tubulin/histoneH2B-mRFP*) mitosis were used for analysis. Cells were bleached along the equator of the mitotic spindle shortly after NEBD once the spindle shape was formed, before anaphase entry. 38-pixel width slices covering the length of the spindle were cut out and arranged using the montage tool in FIJI for easy viewing of the movement of the bleached region, where their slopes were then taken as MT flux rate.

#### Protonema filament bend frequency

Epifluorescence images of moss protonema filaments cultured on BCDAT media were used for analysis in FIJI. Contrast was adjusted to make edges of protonema filaments clearer. A blind test was performed to ascertain waviness threshold, where acute angles above 18° were determined to be ‘wavy’. Randomly chosen protonema filaments were analysed for all the angles of bends along a length of protonema filament using the FIJI in-built angle tool. The number of bends that were ‘wavy’ (>18°) were counted, and taken over the length measured as bend frequency.

#### MT foci trajectories

The MT foci of *Kinesin-13* KO, *-8* KO, and control moss expressing GFP-tubulin were imaged at 60x magnification with z-series every 0.3 µm for a range of 20 µm every 3 min for 3 h. Maximum z-projection of the files were analysed using FIJI MOSAIC plug-in (Sbalzarini and Koumoutsakos, 2005) particle tracker 2D/3D (radius: 20, cut-off: 0.001, per/abs: 0.005, link range: 5, displacement: 20, dynamics: Brownian) to automatically generate the MT foci trajectories. The linear regression of the trajectories was rotated to horizontal and normalised to start at (x, y) = (0, 0) in order to plot graphs in Figure 5D.

#### Interphase endoplasmic MT plus-end dynamics

For plus-end shrink rate, catastrophe frequency, and rescue frequency, oblique illumination time-lapse images taken every 3 s of the interphase endoplasmic MT network of *Kinesin-13* KO (GPH0438#30) and control (*GFP-tubulin/histoneH2B-mRFP*) were analysed. For MT shrink rate, kymographs were drawn on five randomly chosen MTs per cell (cell N = 25; total MT N = 125), and the slope was taken as shrink rate. To analyse catastrophe and rescue frequencies, ∼5 x 6 µm area in a cell was randomly selected whereupon every traceable MT plus-end was traced for a kymograph to count the number of catastrophe or rescue events over the observed growth or shrink time respectively. For ease of tracking, areas with fewer MTs but were not near the cell plate were preferred. Two independent experiments were performed and consistent. To determine MT growth rate, *Kinesin-13* KO moss expressing EB1-Citrine was used, where Citrine marks growing MT plus-ends. Spinning disc time-lapse images taken every 3 s of the interphase endoplasmic MT array in *Kinesin-13* KO (GPH0577#2, 11) and control (*EB1-Citrine/mCherry-tubulin*, GPH0379#2) moss were analysed. Kymographs were taken along the edge of the apical side of the tip cell of protonema filaments. Slopes of EB1-Citrine comets in kymographs were measured and taken as growth rate.

#### Statistics

Welch’s two-sample t-test (two-sided) was used when samples were approximately normally distributed. Tukey multiple comparison test was used for Figure 5B. Significance with the following p-values are indicated as follows: *: < 0.05; **: < 0.01; ***: < 0.001; ****: < 0.0001.

## Accession numbers

*Physcomitrella patens Kinesin-13a*, Pp3c1_27370; *Kinesin-13b*, Pp3c14_9830; *Kinesin-13c*, Pp3c10_106980; *Kinesin-8Ia*, Pp3c1_32120; *Kinesin-8Ib*, Pp3c2_3070; *Kinesin-8II*, Pp3c21_9290.

## Author contributions

MY and GG conceived project; SYL and MY designed and performed experiments; SYL, TE, and MY analysed experimental data; TE performed computer simulation; SYL, GG, and MY wrote paper.

## Acknowledgements

We would like to thank Yoshihisa Oda, Momoko Nishina and Tomohiro Miki for moss lines and plasmids; Yoshikatsu Sato and Advanced Imaging Support (ABiS) platform (JP16H06280) for help of photobleaching experiments; Elena Kozgunova, and Rie Inaba for technical assistance, Mitsuyasu Hasebe for reagents, and Shogo Takatani for discussions. This work was funded by JSPS KAKENHI (17H06471 to G.G. and 19K23723 to M.Y.) and by JSPS and DFG under the Joint Research Projects-LEAD with UKRI (to G.G.). The authors declare no competing interests.

## Movie legends

Movie 1. Mitosis of control, Kinesin-13, and Kinesin-8 KO moss

GFP-tubulin and histoneH2B-mRFP were imaged with spinning disc confocal microscopy. NEBD, 0 min; Playback at 10 fps at 1 min intervals; left, basal side; right, apical side.

Movie 2. Spindle poleward flux of control and Kinesin-13 KO moss

GFP-tubulin on the mitotic spindle at metaphase was photobleached in a strip (0 min), and the migration of the photobleached strip towards the poles can be observed. Playback at 20 fps at 3 s intervals.

Movie 3. Protonema filament growth of control, Kinesin-13 and -8 KO moss.

GFP-tubulin was imaged with epifluorescence microscopy. Playback at 30 fps at 3 min intervals.

Movie 4. MT foci in protonema growth of control, Kinesin-13 and -8 KO moss

The MT foci was imaged with spinning disc confocal microscopy. Movies are maximum z-projections of z-stacks taken every 0.3 µm for a 20 µm range. Playback at 15 fps at 3 min intervals.

Movie 5. Localisation of Kinesin-13 during interphase

Kinesin-13c is shown representatively for the other two paralogues, which show similar localisation. Localisation at the MT foci (top panel) and the interphase endoplasmic MT array (bottom panel) were imaged with spinning disc confocal microscopy and oblique illumination fluorescence microscopy respectively. Playback at 15 fps at 3 s intervals.

